# Zonation-dependent single-endothelial cell transcriptomic changes in the aged brain

**DOI:** 10.1101/800318

**Authors:** Lei Zhao, Zhongqi Li, Joaquim S. L. Vong, Xinyi Chen, Hei-Ming Lai, Leo Y. C. Yan, Junzhe Huang, Samuel K. H. Sy, Xiaoyu Tian, Yu Huang, Ho Yin Edwin Chan, Hon-Cheong So, Wai-Lung Ng, Yamei Tang, Wei-Jye Lin, Vincent C.T. Mok, Ho Ko

## Abstract

With advances in single-cell genomics, molecular signatures of cells comprising the brain vasculature are revealed in unprecedented detail^1,2^, yet the ageing-associated cell subtype transcriptomic changes which may contribute to neurovascular dysfunction in neurodegenerative diseases^3–7^ remain elusive. Here, we performed single-cell transcriptomic profiling of brain endothelial cells (EC) in young adult and aged mice to characterize their ageing-associated genome-wide expression changes. We identified zonation-dependent transcriptomic changes in aged brain EC subtypes, with capillary ECs exhibiting the most transcriptomic alterations. Pathway enrichment analysis revealed altered immune/cytokine signaling in ECs of all vascular segments, while functional changes impacting the blood-brain barrier (BBB) and glucose/energy metabolism were most prominently implicated in ECs of the capillary bed – the primary site where ECs and other neurovascular unit (NVU) cell types closely interact and coordinate to regulate BBB and cerebral blood flow in health and diseased conditions^8–17^. Furthermore, an overrepresentation of Alzheimer’s disease (AD)-associated genes identified from GWAS studies was evident among the human orthologs of differentially expressed genes of aged capillary ECs but not other EC subtypes. Importantly, for numerous EC-enriched differentially expressed genes with important functional roles at the BBB and/or association with AD, we found concordant expression changes in human aged or AD brains. Finally, we demonstrated that treatment with exenatide, a glucagon-like peptide-1 receptor (GLP-1R) agonist, strongly reverses transcriptomic changes in ECs and largely reduces BBB leakage in the aged brain. Thus, our study provides insights into detailed transcriptomic alterations underlying brain EC ageing that are complex with subtype specificity yet amenable to pharmacological interventions.

## Introduction

Ageing is the strongest risk factor for neurodegenerative diseases. During the ageing process, complex cellular and functional changes occur in the brain. These include neurovascular dysfunction manifesting in blood-brain barrier (BBB) breakdown associated with microscopic pathological changes of vascular cells^9,18–20^. Forming the innermost lining of the vasculature, brain endothelial cells (ECs) closely interact with other cell types of the neurovascular unit (NVU) to regulate diverse functions including BBB integrity^8,12,13^, immune signaling and brain metabolism, all of which are altered in the aged brain^18,21–23^. These imply that ECs play crucial roles in the brain ageing process that are still incompletely understood. With advances in single-cell genomics, especially single-cell RNA sequencing (scRNA-seq), the molecular subtypes of brain ECs that segregate along the arteriovenous axis have been identified in exquisite details^1,2^. Given the heterogeneity of EC subtypes, the genome-wide expression changes in ageing brain ECs are likely highly complex and yet to be fully revealed.

Uncovering brain EC transcriptomic changes is of paramount importance for enhancing our understanding of neurovascular dysfunction in ageing, which is increasingly recognized as a major contributor to pathogenesis in multiple neurodegenerative diseases^3–7^. In Alzheimer’s disease (AD), postmortem examination revealed that vascular pathology including lacunes, microinfarcts and other microvascular lesions coexist with classical AD pathology (i.e. amyloid plaques and neurofibrillary tangles) in the majority (~80%) of patients^24^. BBB breakdown is an early and consistent feature preceding symptom onset in AD patients^3,6,19,25,26^. In multiple transgenic animal models of AD harboring amyloid precursor protein mutations and/or ApoE4, BBB leakage also precedes and accelerates the development of parenchymal amyloid deposition, neurofibrillary tangles formation, neuronal dysfunction and behavioral deficits^27^. In numerous other neurodegenerative diseases, such as frontotemporal dementia (FTD), amyotrophic lateral sclerosis (ALS) and Parkinson’s disease (PD), functional and pathological microvascular changes occur in respective disease-affected brain regions^4,7^. While disease-specific factors differentially contribute to neurovascular defects, ageing-associated neurovascular dysfunction represents a shared pathological component among the distinct neurodegenerative diseases. It remains to be determined, how ageing differentially influences different molecular subtypes of brain ECs, resulting in neurovascular dysfunction and neurodegenerative disease susceptibility.

Moreover, proving the reversibility of ageing-associated transcriptomic and functional changes, preferably by pharmacological means, would bear significant implications for the development of preventive or disease-modifying therapeutics. Originally developed to potentiate glucose-induced insulin release and treatment of diabetes, glucagon-like 1-peptide receptor (GLP-1R) agonists were reported to have therapeutic efficacy in multiple animal models of neurodegenerative diseases including AD, ALS, PD^28–33^ and in human trials of AD and PD^34,35^. In the brain, GLP-1R is expressed by multiple cell types, including neurons, microglia and potentially vascular cells^32,36–38^. It remains to be tested, whether GLP-1R agonists may act partly via reversal of neurovascular ageing, accounting for the general applicability as potential therapeutics for multiple neurodegenerative conditions.

To gain insights into the complex landscape of gene expression changes in aged brain vasculature, we compared single-cell transcriptomes of brain ECs from young adult and aged mouse brains, employing arteriovenous zonation markers to classify EC subtypes. We seek to address several core questions, including: (1) what common and specific gene expression changes occur in different EC subtypes along the arteriovenous axis in the aged brain, (2) whether genes related to BBB functionality and neurodegenerative conditions in aged brain ECs exhibit arteriovenous zonation-specific differential expression, (3) whether some transcriptomic alterations in aged mouse brain ECs may generalize to human aged or AD brains, and (4) if the ageing-associated transcriptomic changes of brain ECs are amenable to pharmacological intervention by GLP-1R agonist treatment with accompanying improvement in BBB integrity.

## Results

### scRNA-seq identifies zonation-dependent endothelial cell transcriptomic changes in aged mouse brain

To obtain ageing-associated genome-wide expression changes of neurovascular cells, we performed high-throughput scRNA-seq of brain vascular cells from young adult (2 – 3 months old) and aged (18 – 20 months old) C57BL/6J mice (*n* = 5 mice for each group) (Fig. 1a). After quality control filtering (see Methods), we obtained a total of 43922 single-cell transcriptomes. These included 31555 cells from young adult and 12367 cells from aged mouse brains (Fig. 1a), among which there were 12357 brain ECs, 1425 smooth muscle cells (SMC), 642 pericytes (PC), 9642 astrocytes (AC) and 11767 microglia (MG), alongside numerous other major brain cell type clusters (Fig. 1b, Supplementary Fig. 1a). We employed a probabilistic method^39^ (see Methods) to assign each of the ECs from young adult (*n* = 8627 cells) and aged (*n* = 3730 cells) mouse brains to one of six molecular subtypes with distinct arteriovenous distributions based on the expression profiles of 267 arteriovenous zonation-dependent genes in ECs^1^ (Fig. 1c, Supplementary Fig. 1b, also see Supplementary Table 1 for lists of the genes). These correspond to two putative arterial subtypes (aEC1, *n* = 1141 cells and 526 cells; aEC2, *n* = 998 and 679 cells from young adult and aged groups respectively), capillary subtype (capEC, *n* = 1587 and 697 cells), venous subtype (vEC, *n* = 798 and 412 cells), and two subtypes with mixed distributions (arterial/venous – avEC, *n* = 293 and 229 cells; and capillary-venous EC – vcapEC, *n* = 3122 and 801 cells) (Fig. 1c). For both age groups, the relative expression levels of arteriovenous zonation marker genes including those of artery/arteriole (Bmx and Vegfc), capillary (Mfsd2a and Tfrc), vein (Nr2f2 and Slc38a5), and artery/vein (Vwf and Vcam1) match the respective spatial designations (Fig. 1d, Supplementary Fig. 1c). Across the two age groups, the proportions of capEC subtype were similar (around 20% for both groups), while we obtained more vcapEC from the young than aged adult mouse brains (39.3% and 24.0% respectively, Fig. 1e).

**Fig. 1.**
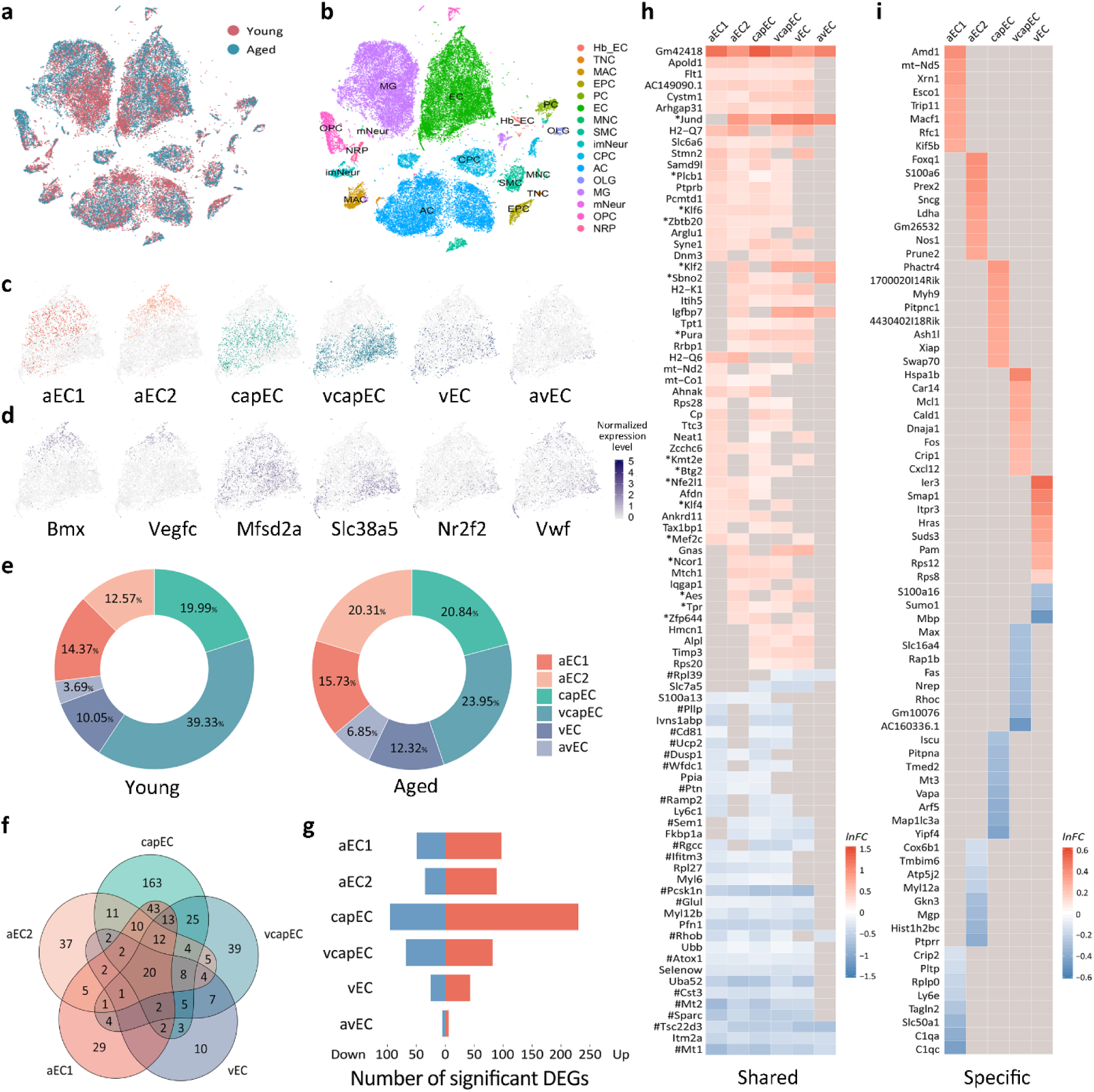
**a,** t-SNE visualization of single-cell transcriptomes from young adult (2 – 3 months old, *n* = 31555 cells from 5 mice, brown) and aged (18 – 20 months old, *n* = 12367 cells from 5 mice, blue) mouse brains. **b,** Primary cell type clusters identified based on marker gene expression patterns. Cell type abbreviations: EC, endothelial cell; SMC, smooth muscle cell; PC, pericyte; MG, microglia; AC, astrocyte; NRP, neuronal restricted precursor; OPC, oligodendrocyte precursor cell; OLG, oligodendrocyte; mNeur, mature neuron; imNeur, immature neuron; EPC, ependymocyte; CPC, choroid plexus epithelial cell; Hb_EC, hemoglobin-expressing vascular cell; MAC, macrophage; TNC, tanycyte; MNC, monocyte. **c,** Endothelial cell (EC) subtypes, including two arterial (aEC1, *n* = 1141 cells and 526 cells; aEC2, *n* = 998 and 679 cells from young adult and aged groups respectively), capillary (capEC, *n* = 1587 and 697 cells), venous and capillary (vcapEC, *n* = 3122 and 801 cells), venous (vEC, *n* = 798 and 412 cells) and arterial-venous (avEC, *n* = 293 and 229 cells) subtypes. Different colors are used for the subtypes to facilitate visualization. **d,** Expression patterns of arterial (Bmx, Vegfc), capillary (Mfsd2a), venous (Slc38a5, Nr2f2) and arterial/venous (Vwf) marker genes in ECs visualized by t-SNE. 6000 cells were subsampled and shown for better clarity of visualization. **e,** Proportions of EC subtypes obtained from each age group. Scale bar: normalized expression level. **f,** Venn diagram showing the overlap of significant differentially expressed genes (DEGs) (FDR adjusted *P*-value < 0.05 and |*lnFC*| > 0.1) between five EC subtypes. **g,** Numbers of significant upregulated (red bars) and downregulated (blue bars) DEGs for each EC subtype. **h,** Heatmap of shared (significant in ≥ 3 EC subtypes) upregulated (upper panel) and downregulated (lower panel) DEGs in the aged brain. Remarks: *Transcription factor / regulatory genes and #stress response genes by the Gene Ontology Annotation. **i,** Heatmap of EC subtype-specific upregulated (upper panel) and downregulated (lower panel) DEGs (i.e. significant differential expression with adjusted *P*-value < 0.05 and |*lnFC*| > 0.1 in only one EC subtype). Up to eight are shown for each EC subtype.

To assay how ageing may commonly and differentially influence brain EC subtype gene expression in different segments of the vascular network, we calculated differentially expressed genes (DEGs) expressed in natural log(fold change) (*lnFC*) in aged relative to young adult mouse group for each of the EC subtypes (Fig. 1f-i). DEGs were considered significant only if reaching statistical significance (FDR-adjusted *P*-value < 0.05) with |*lnFC*| > 0.1. Among the EC subtypes, aged capEC had the largest number of significant DEGs while avEC had the least expression changes (capEC: 325; vcapEC: 150; aEC1: 146; aEC2: 124; vEC: 68; avEC: 11; Fig. 1f, g). For all EC subtypes, we found more upregulated than downregulated DEGs (Fig. 1g). Altogether, the EC subtype significant DEGs constituted a total of 469 unique genes. 21.96% (103 out of 469) of the DEGs were not significant for all EC pooled (Supplementary Fig. 2a), while those significant tend to have magnitudes of expression changes underestimated (Supplementary Fig. 2b). These highlighted the importance of fine subtype classification in addition to the localization of expression changes with respect to zonation. 192 significant DEGs were shared by more than one EC subtype, almost all (191 out of 192, 99.5%) had conserved directionality of change across different EC subtypes.

A set of 55 shared genes upregulated in multiple EC subtypes (≥ 3 subtypes) were identified, among which only 7 were significant in at least five EC subtypes (Fig. 1h). Among the shared upregulated DEGs there was an abundance of transcription factor or regulatory genes (e.g. Btg2, Klf2, Ncor1, Sbno2, Zbtb20) (Fig. 1h). A smaller set of 34 commonly downregulated genes (≥ 3 EC subtypes) was identified, among which only 15 were shared by at least five aged brain EC subtypes (Fig. 1h). Several of the shared downregulated genes (e.g. Cst3, Pcsk1n and Sparc) were previously reported to be generally downregulated in EC and other aged brain cell types^40^ (Fig. 1h). Numerous stress response genes (e.g. Atox1, metallothionein genes Mt1 and Mt2, Sem1, Tsc22d3) were present amongst the commonly downregulated genes (Fig. 1h). A total of 277 significant DEGs were specific to only one EC subtype (194 upregulated and 83 downregulated), with 163 being capEC DEGs (120 upregulated and 43 downregulated; Fig. 1f, i) and the rest specific to one of the other EC subtypes except avEC (Fig. 1f, i). Brain ECs therefore have both shared and divergent ageing-associated differential expressions, with the most prominent changes occurring in capillary ECs.

### Zonation-dependent functional changes implicated in aged brain EC transcriptomes

Age-dependent endothelial dysfunction is a major contributor to neurovascular dysfunction. To identify altered functional pathways implicated in the aged brain EC expression changes, we next performed pathway analysis on significant DEGs of each EC subtype (except avEC due to a low number of significant DEGs) (Fig. 2a). Among capEC DEGs, we found significant enrichment of genes associated with adherens junction (AJ) (e.g. Afdn, Ctnna1, Iqgap1, Lef1, Ptprb) and tight junction (TJ) (e.g. Arhgef2, Cgnl1, Nedd4, Ocln, Ppp2r2a) (Fig. 2a, b). Prompted by this observation, we further examined patterns of expression changes of AJ, TJ component-encoding genes, and genes that regulate BBB transcellular transport in different brain EC subtypes. The claudin 5-encoding gene Cldn5 has been reported to be downregulated in ECs in general^40^. In our data, a significant downregulation of Cldn5 occurred only in aged vcapEC (Fig. 2b). We also found downregulated expression of Mfsd2a in aged capEC, which encodes the major facilitator superfamily domain-containing protein 2 (MFSD2A, or sodium-dependent lysophosphatidylcholine symporter 1) and is crucial for limiting transcytosis in the BBB^41^ (Fig. 2b).

**Fig. 2.**
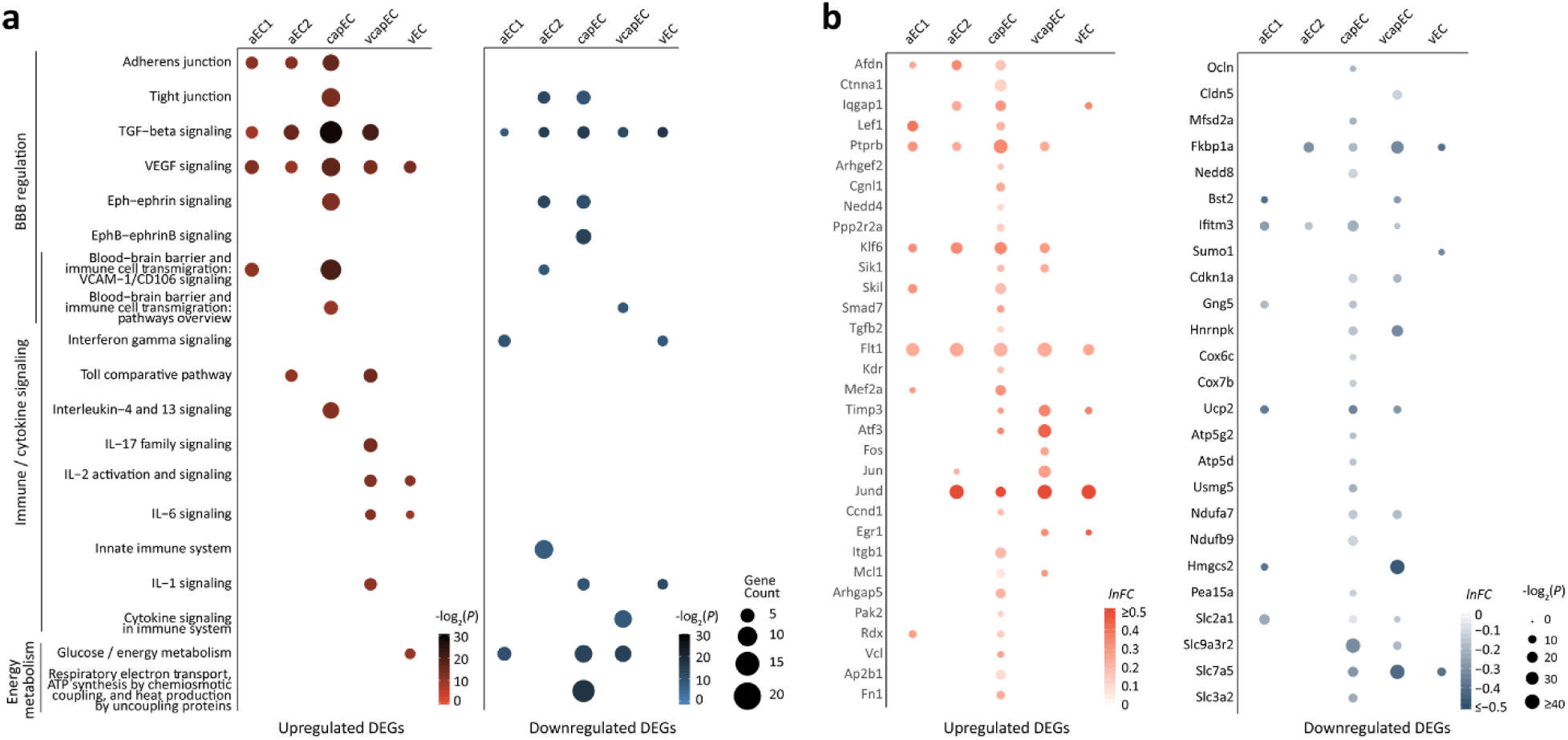
**a,** Dot plots of BBB-regulatory, immune/cytokine signalling, respiratory electron transport chain and glucose/energy metabolism pathways with significant enrichment in the aged brain for upregulated (left panel) and downregulated (right panel) DEGs. **b,** Dot plots showing differential expression profiles of selected upregulated (left panel) and downregulated (right panel) genes in aged brain ECs associated with enriched pathways shown in **a**. in the different EC subtypes.

In addition to AJ- and TJ-associated genes, numerous growth factors, immune and cell-to-cell junction signaling pathways are also important regulators of BBB integrity. TGF-β signaling-related genes were significantly enriched in the DEGs of all five aged EC subtypes analyzed (e.g. upregulated: Klf6, Sik1, Skil, Smad7, Tgfb2; downregulated: Fkbp1a, Nedd8) (Fig. 2a, b). The upregulation of VEGF-signaling pathway genes also impacts brain ECs of all segments (e.g. Flt1, Kdr, Mef2a, Timp3) (Fig. 2a, b). Furthermore, the enrichment of immune/cytokine signaling-related gene sets was observed for all vascular segments. These include genes mediating interferon gamma response in aEC1 and vEC (e.g. downregulated: Bst2, Ifitm3, Sumo1), toll-like receptor signaling in aEC2 and vcapEC (e.g. upregulated: Atf3, Fos, Jun, Jund), as well as interleukin signaling in capEC, vcapEC and vEC (e.g. upregulated: Ccnd1, Egr1, Itgb1, Mcl1; downregulated: Cdkn1a). Overall, capEC had the most prominent BBB-regulatory signaling pathways enrichment, which also included VCAM1-mediated immune cell transmigration (e.g.: upregulated: Arhgap5, Pak2, Rdx, Vcl) and Eph-ephrin/EphB-ephrinB signaling (e.g. upregulated: Ap2b1, Fn1; downregulated: Gng5, Hnrnpk) associated genes (Fig. 2a, b).

Altered energy metabolism may also be a key contributor to brain endothelial ageing^42^. We found that respiratory electron transport chain / ATP synthesis (e.g. cytochrome oxidase subunit genes Cox6c and Cox7b, Ucp2, multiple ATP synthase and NADH:ubiquinone oxidoreductase genes) and glucose/energy metabolism (e.g. Hmgcs2, Pea15a and multiple SLC transporter genes) associated genes were most enriched in the downregulated DEGs of aged capEC, and to a lesser extent in aEC1 and vcapEC (Fig. 2a, b). Of note, Slc2a1 (encoding the glucose transporter GLUT1) was found downregulated in multiple EC subtypes (Fig. 2b). Therefore, while impaired energy metabolism has been most strongly implicated in capEC, other vascular segments are also likely impacted in the aged brain.

Collectively, these results revealed a complex pattern of ageing-related transcriptomic changes in specific EC subtypes that may impact BBB function, especially in capEC. Important subsets of the DEGs were related to immune/cytokine signalling and energy metabolism, and may jointly contribute to age-dependent BBB dysfunction and susceptibility to neurodegeneration.

### Aged brain EC transcriptomic changes and neurodegenerative disease association

Genome-wide association studies (GWAS) identified many associated genetic variants of cerebrovascular and neurodegenerative diseases with a vascular component^43–54^. We asked whether some of these genes would have ageing-associated expression changes in ECs (see Supplementary Table 2 for disease-associated gene lists used for this analysis). Indeed, we found a total of 40 genes among the DEGs of aged brain ECs associated with one or more of 6 out of the 11 diseases examined (Fig. 3), 32 of which were specific to ECs with no significant differential expression found in other major brain cell types in our dataset (Fig. 3). Overall, 31 disease-related genes identified were significant DEGs of capEC (Fig. 3), and 23 are AD-associated (see Supplementary Table 3 for disease association summary). Interestingly, the abundance of AD GWAS genes among the DEGs was not merely a result of the large number of AD GWAS genes, as they were significantly overrepresented in aged capEC DEGs (Fig. 3, number of overlapped genes = 18; *P* = 0.022, hypergeometric test). While numerous risk genes of other neurovascular and neurodegenerative diseases also overlapped with aged EC DEGs, they did not reach statistical significance for overrepresentation in any EC subtypes (Fig. 3, e.g. number of overlapped genes for stroke genes and capEC DEGs = 4; *P* = 0.067). Notably, we found upregulation of Cd2ap (Fig. 3), which has previously shown to be required for the maintenance of normal BBB^55^ and its loss-of-function variants may contribute to AD pathogenesis by impairing BBB integrity. We also noted upregulation of Dlc1 in aged capEC (Fig. 3), which has also been identified as an upregulated gene in human AD patient brain ECs^56^. Our data thus support that AD has a unique vascular ageing component, especially in the capillary endothelium, among other neurodegenerative diseases.

**Fig. 3.**
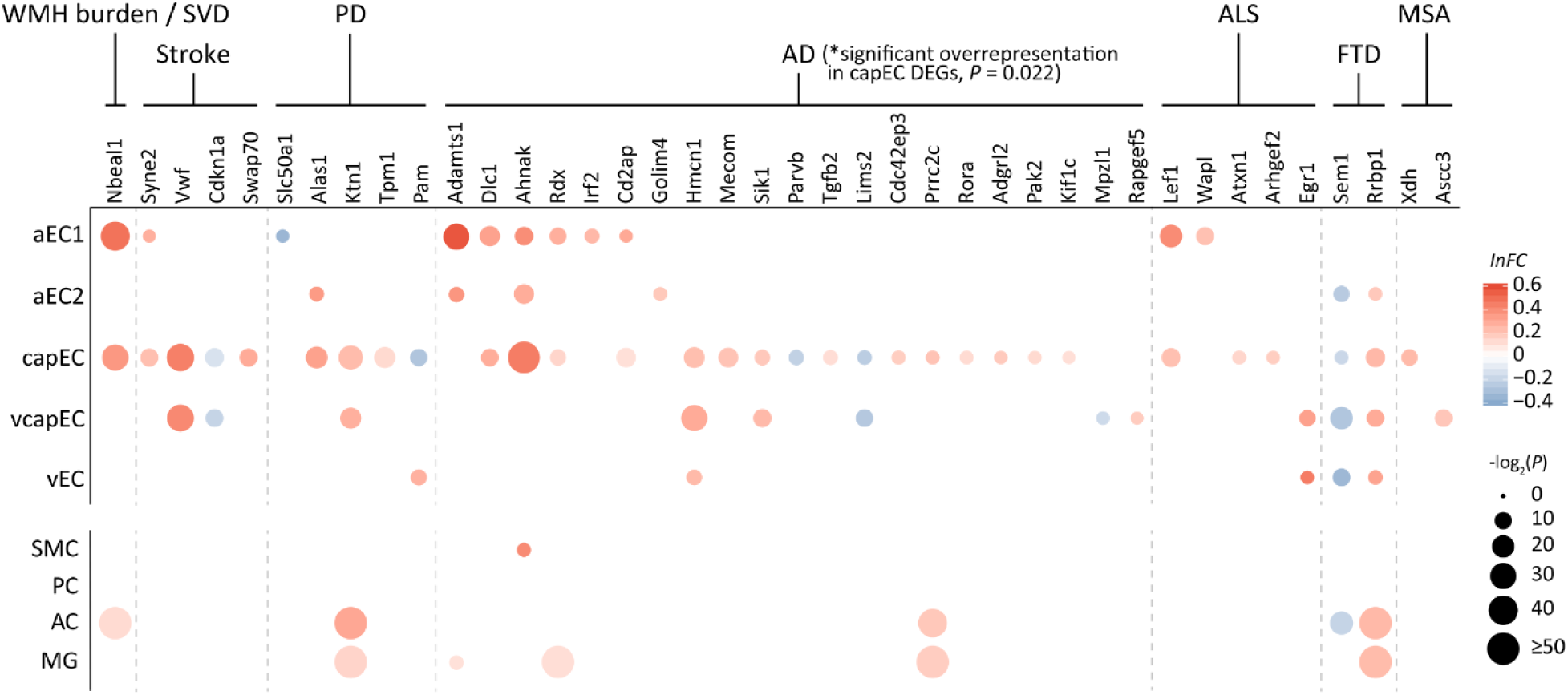
Differential expression profiles of aged brain EC subtype DEGs whose human orthologs are GWAS genes of cerebrovascular or neurodegenerative diseases examined. capEC DEGs had the most overlap with a significant overrepresentation of AD-GWAS genes (*P* = 0.022, hypergeometric test). The differential expressions of these genes in vascular mural cells (SMC: smooth muscle cell; PC: pericyte), astrocyte (AC) and microglia (MG) are also shown. Disease abbreviations: WMH burden: white matter hyperintensity burden; SVD: cerebral small vessel disease; PD: Parkinson’s disease; AD: Alzheimer’s disease; ALS: amyotrophic lateral sclerosis; FTD: frontotemporal dementia; MSA: multiple systems atrophy.

### Concordant and discordant EC transcriptomic changes in normal aged and AD human brains

We speculated that some of the transcriptomic changes found in aged mouse brain ECs are conserved and detectable in normal human aged brains with bulk RNA sequencing data, available from the Genotype-Tissue Expression (GTEx) project database^57^. We divided the data in the database into two age groups (≥ 60 versus < 60 years old) and identified aged mouse brain EC subtype DEGs whose human orthologs had significant differential expression in aged human brains. The human orthologs of aged mouse brain EC-enriched DEGs had a much higher proportion of significant differential expression than all genes profiled (38.1%, 40 out of 105 EC-enriched genes vs 6.5%, 3424 out of 52669, *P* < 1 × 10^−5^; Fig. 4a) (see Methods). Of note, the majority of these DEGs were upregulated in aged human brains (92.5%, 37 out of 40). Manual review of the Human Protein Atlas^58^, Allen Brain Atlas human brain single nucleus RNA-seq data^59^ and available literature confirmed human brain EC expression for most of these genes (95%, 38 out of 40, see Supplementary Table 4). 20 human homologs of upregulated aged mouse brain EC DEGs had concordant increase in expression in aged human brains (Fig. 4a, b; Supplementary Fig. 3a). These included numerous BBB regulation-associated genes (e.g. Iqgap1, Ptprb, Timp3, Flt1). Multiple DEGs with concordant changes are also AD-associated genes (including P-glycoprotein gene Abcb1a, Adamts1, Ahnak, Mecom) (Fig. 4a, b). Notably, P-glycoprotein (P-gp, human ortholog: ABCB1), exclusively expressed by ECs in human brain (Supplementary Table 4), mediates brain-to-blood amyloid beta clearance and its diminished function is thought to contribute to AD^60,61^. The increased expression of P-gp in aged brains may represent an adaptive change. However, 19 EC-enriched genes had different directionality of change in human aged brains, among which most (89.5%, 17 out of 19) had increased expression in aged human brains but downregulated in aged mouse brain ECs (Fig. 4a). These included several genes with important functions at the BBB, such as Cldn5, Slc2a1, Ifitm3 (≥ 60 versus < 60 years old human brain, FDR-adjusted *P*-value < 0.05) (Fig. 4a, b). Additionally, Mfsd2a exhibited a trend of upregulation in aged human brains (≥ 60 versus < 60 years old human brain, FDR-adjusted *P*-value = 0.0505) (Fig. 4b).

**Fig. 4.**
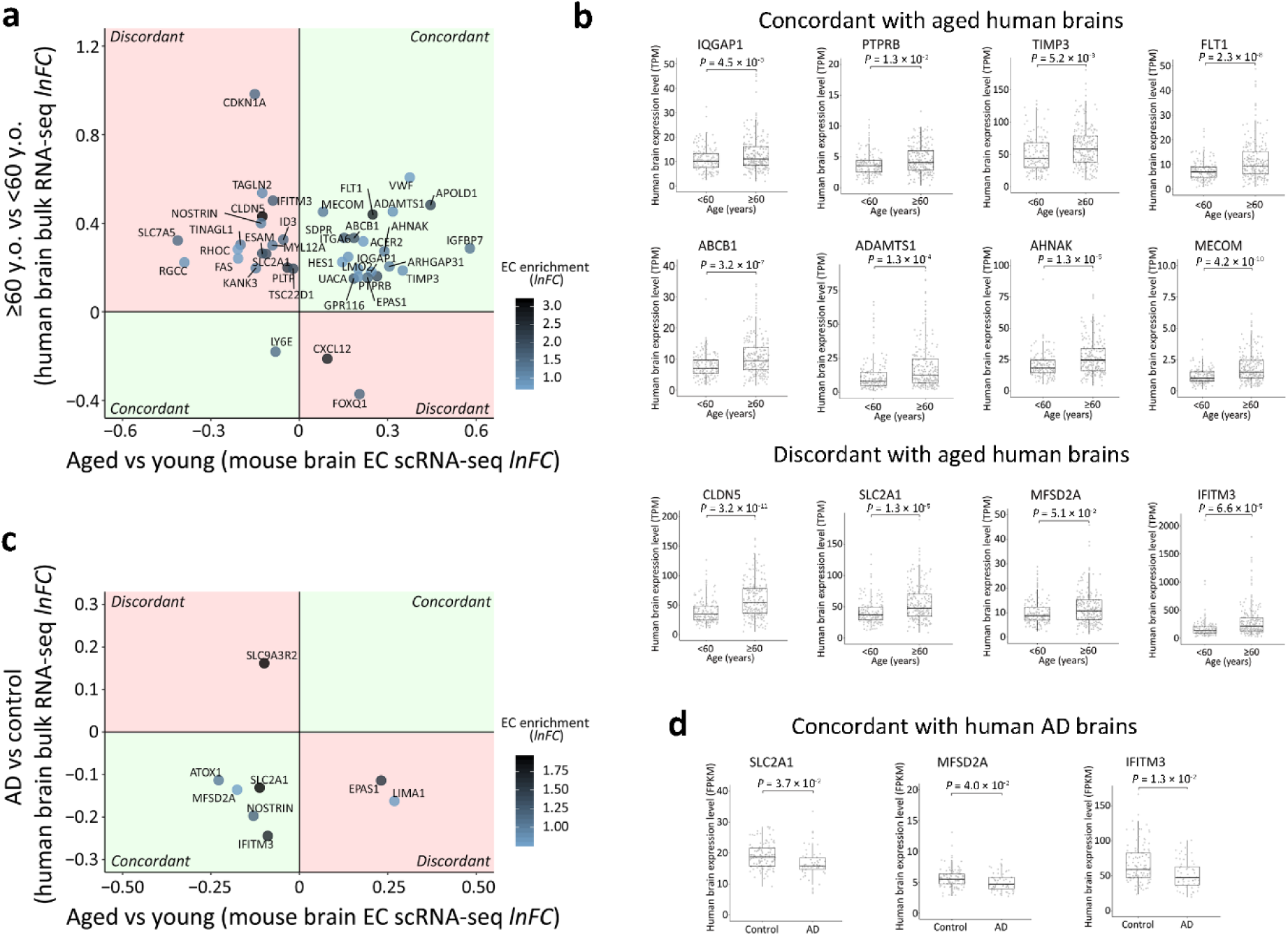
**a,** Aged human brain (≥ 60 years old) differential expressions relative to younger subjects (< 60 years old) (FDR-adjusted *P*-value < 0.05, y-axis) plotted against expression changes in pooled aged mouse brain ECs (x-axis), showing aged mouse brain EC subtype DEGs whose human orthologs had concordant (first and third quadrants, light green) or discordant (second and fourth quadrants, light red) expression changes in human AD brains. Only DEGs with at least two-fold enrichment in EC (i.e. *lnFC* of expression in ECs relative to other cell types > 0.7) were included, with color of dots representing the degree of EC enrichment. **b,** Human brain expression levels of selected genes associated with BBB regulation (Iqgap1, Ptprb, Timp3, Flt1) and AD (P-gp gene, Adamts1, Ahnak, Mecom) with concordant changes in aged human and mouse brains (≥ 60 versus < 60 years old human brains, FDR-adjusted *P*-value < 0.05), or with important functional roles at the BBB and yet discordant changes (≥ 60 versus < 60 years old human brains, Cldn5, Slc2a1 and Ifitm3: FDR-adjusted *P*-value < 0.05; Mfsd2a: FDR-adjusted *P*-value = 0.0505). **c,** Human AD brain differential expressions relative to age-matched control subjects (FDR-adjusted *P*-value < 0.05, y-axis) plotted against expression changes in pooled aged mouse brain ECs (x-axis), showing aged mouse brain EC DEGs whose human orthologs had concordant (first and third quadrants, light green) or discordant (second and fourth quadrants, light red) expression changes in human AD brains. Similar to **a,** only DEGs with at least two-fold enrichment (*lnFC* of EC expression relative to other cell types > 0.7) are shown, with color of dots representing the degree of EC enrichment. **d,** Human brain expression levels of selected genes with important functional significance and concordant expression changes in human AD relative to control subjects and aged mouse brains (AD versus control human brains, Slc2a1, Mfsd2a and Ifitm3: FDR-adjusted *P*-value < 0.05).

We then attempted to further identify aged mouse brain EC subtype DEGs with the same expression changes in human AD brains. We calculated DEGs for human AD versus control brains with no dementia (aged 78 – over 100 years old), from the Allen Brain Institute Aging, Dementia and Traumatic Brain Injury study dataset of human brain bulk RNA sequencing^62,63^. After selecting for EC-enriched genes, the directionality of change of the human AD DEGs were compared to the EC subtype DEGs. This revealed 5 EC-enriched genes with the same changes found in aged mouse and human AD patient brains, 4 of which had evidence of expression in human brain ECs (Supplementary Table 4). In stark contrast to those DEGs with matching altered expression in normal aged mouse and human brains, all 5 genes had reduced expression in human AD brains (Fig. 4c; Supplementary Fig. 3b). Interestingly, several of these genes with important functional roles at the BBB had concordant downregulation in aged mouse and AD brains (Fig. 4c, d), despite discordant expression changes in aged mouse and human brains (Fig. 4a, b). For instance, consistent with previous reports, reduced expression of Slc2a1 in AD brains was found^64–67^ (Fig. 4c, d). On the other hand, the aged mouse brain capEC-downregulated DEG Mfsd2a (Fig. 2b) also had reduced expression in AD brains compared to non-demented aged human brains, suggesting a possible contribution to BBB breakdown in AD (Fig. 4c, d). Ifitm3, an IFN-responsive gene that plays an important role in restricting viral invasion^68–70^ and upregulated in normal aged human brains, was also found downregulated in multiple aged mouse brain EC subtypes (Fig. 2b) and human AD brains (Fig. 4c, d). Therefore, in addition to the overrepresentation of AD GWAS genes, we have found a multitude of similar expression changes in the normal aged mouse and human AD brains that are functionally important and may contribute to degenerative changes.

### GLP-1R agonist treatment reverses ageing-associated EC transcriptomic changes and reduces BBB leakage

Finally, we asked if the complex cell subtype-dependent transcriptomic alterations of aged vascular cells are amenable to pharmacological intervention with improvements in NVU integrity. Compared to young adult (2 – 3 months old) mice, aged (18 – 20 months old) mice showed nearly 15.8-fold increase of 40 kDa TRITC-conjugated dextran (TRITC-dextran) extravasation in the cerebral cortex (see Methods), signifying increased BBB leakage (Fig. 5a, b; volume of extravasated TRITC-dextran in aged vs young adult group, *P* < 0.001, *n* = 9 image stacks from 3 mice for each group; see Methods). Strikingly, 4 – 5 week treatment of aged mice by the GLP-1R agonist exenatide (5 nmol/kg/day, intraperitoneal injection starting at 17 – 18 months old) reduced TRITC-dextran extravasations in the cerebral cortex by more than 50% (Fig. 5a, b, *P* < 0.001 compared to 18 m.o. untreated group, *n* = 9 image stacks from 3 mice for each group).

**Fig. 5.**
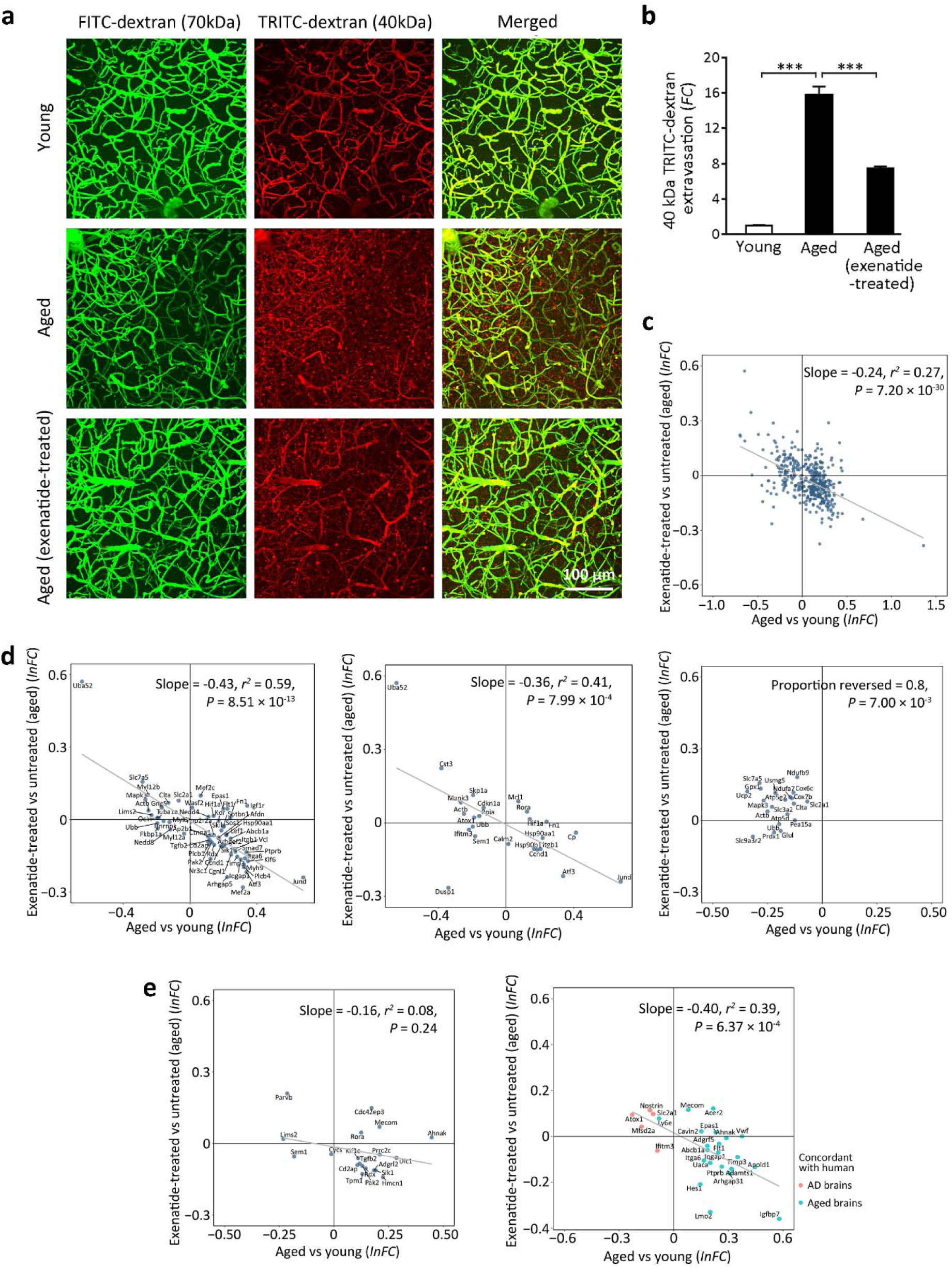
**a,** Three-dimensional rendered images (top view) of *in vivo* two-photon imaging of cerebral vasculature and blood-brain barrier (BBB) leakage in the mouse somatosensory cortex by co-injection of 70 kDa FITC-conjugated dextran (FITC-dextran, green) and 40 kDa TRITC-conjugated dextran (TRITC-dextran, red). FITC-dextran remained in the vasculature and allowed reconstruction of vessels, while extravasation of TRITC-dextran serves as an indicator of BBB leakage which was quantified for young adult, aged and exenatide-treated aged mouse groups. **b,** Volumetric quantification of TRITC-dextran extravasation showing BBB breakdown in aged (18 – 20 months old) relative to young adult mice (2 – 3 months old) (mean fold change in volume of extravasated TRITC-dextran relative to young adult group ± S.E.M. = 15.8 ± 0.9; \*\*\**P* < 0.001 for aged vs young adult mouse group), which was significantly reduced by exenatide treatment (5 nmol/kg/day I.P. for 4 – 5 weeks starting at 17 – 18 months old, mean fold change relative to young adult group ± S.E.M. = 7.5 ± 0.2; \*\*\**P* < 0.001 for exenatide-treated vs untreated aged mouse group). **c-d,** Reversal of brain capillary EC overall (**c**), BBB-regulatory (**d**, left panel), immune/cytokine signalling (**d**, middle panel) and energy metabolism (**d**, right panel) associated gene expression changes by exenatide treatment in aged mouse. **e,** Reversal of aged mouse brain EC differential expressions whose human orthologs are AD-GWAS genes in capEC (left panel), or had concordant changes in normal human normal aged or AD brains in all ECs pooled (right panel).

Remarkably, the observed improvement of BBB integrity was associated with reversal of ageing-associated gene expression changes in brain ECs (Fig. 5c-e, Supplementary Fig. 4). For statistically significant capEC DEGs (i.e. adjusted *P*-value < 0.05 regardless of |*lnFC*|), exenatide treatment reversed the expression changes of 76.4% upregulated (217 out of 284, *P* < 1 × 10^−4^ compared to chance level) and 63.4% downregulated DEGs (85 out of 134, *P* = 5.9 × 10^−3^ compared to chance level) (Fig. 5c, *r*^*2*^ = 0.27; *P* = 7.2 × 10^−30^). This reversal effect also applies to other EC subtypes (Supplementary Fig. 4). Classifying the DEGs by functionality, strong reversal was found for BBB-regulatory (Fig. 5d, left panel, 76.7% of DEGs reversed, *r*^*2*^ = 0.59; *P* = 8.51 × 10^−13^) and immune/cytokine signalling (Fig. 5d, middle panel, 66.7% of DEGs reversed, *r*^*2*^ = 0.41; *P* = 7.99 × 10^−4^) related genes identified from pathway analysis in capEC. The majority of downregulated respiratory electron transport chain- and glucose/energy metabolism-associated genes were upregulated by exenatide treatment in capEC (Fig. 5d, right panel, 80% upregulated, *P* = 0.007 compared to chance level). Finally, the observation also generalized to the subcategory of DEGs whose human orthologs are AD-associated (Fig. 5e, left panel, 68.4% of DEGs reversed, *r*^*2*^ = 0.0803; *P* = 0.24) in capEC, or with concordant changes in the normal aged or AD human brains (Fig. 5e, right panel, 65% of DEGs reversed, *r*^*2*^ = 0.391; *P* = 6.37 × 10^−4^) in all ECs pooled. We have thus shown that despite the complexity, the transcriptomic and functional changes in aged brain ECs are amenable to pharmacological interventions.

## Discussions

Advances in single-cell transcriptomic profiling has led to the introduction of the vascular zonation concept, which enables the fine classification of EC subtypes along the arteriovenous axis based on gene expression differences, highlighting their potential functional distinctions^1^. This opens up the possibility of studying how distinct EC subtypes may be selectively vulnerable in neurovascular and neurodegenerative diseases^7^. In this study, we focus on uncovering transcriptomic changes in aged brain ECs subtypes that may underlie neurovascular dysfunction and increased susceptibility to ageing-associated neurodegenerative conditions. This could potentially advance our understanding of normal ageing changes as a common factor in these diseases, paving the way for further studies of how disease-specific factors lead to differentially compromised NVU function in different conditions.

By classifying EC transcriptomes into six major subtypes according to the reported expression patterns of arteriovenous-variable genes, we uncovered both common and specific ageing-associated transcriptomic changes in the different EC subtypes. Among BBB-associated pathways with enrichment of expression changes, altered TGF-β, VEGF and immune/cytokine signalling appear to impact all vascular segments, while others exhibit subtype-specific changes that notably all impact capEC. Despite the partially overlapping zonation of capEC and vcapEC in capillaries, ageing-related expression changes are more prominent in capEC. Although energy metabolism-related pathway genes had altered expression in both arterial and capillary ECs, joint-downregulation of multiple cytochrome oxidase subunit, ATP synthase, NADH:ubiquinone oxidoreductase and SLC transporter genes only occur in capillary ECs. Our findings thus revealed that the fine zonation classification of ECs is directly relevant to their gene expression and functional changes in the aged brain.

To bridge findings from mouse to human, we performed comparative analyses of mouse brain single-cell and human brain bulk RNA sequencing data. For the aged EC DEGs, selection for EC-enriched genes enabled us to identify a subset of genes with concordant expression changes, for which the majority had confirmed expression in the human brain endothelium. Among the matched genes, numerous have functional significance related to BBB regulatory pathways. Detectable differential expressions of these genes from the bulk RNA sequencing data suggest that they are prominent changes in human aged brains. More detailed single-cell transcriptomic studies on human brain ECs are warranted to uncover transcriptomic changes in the ageing brain vasculature and whether a similar arteriovenous zonation principle applies.

In AD, neurovascular dysfunctions manifesting in reduced cerebral blood flow, impaired neurovascular coupling and BBB breakdown precede neurodegeneration and atrophic changes^3,4^. Interestingly, we found an overrepresentation of AD GWAS genes among the human orthologs of the aged capillary ECs DEGs. For a subset of these genes, we also found concordant upregulation of their orthologs in the aged human brain (i.e. P-gp gene, Adamts1, Ahnak, and Mecom). The upregulation of P-gp expression is in apparent contradiction to the reported diminished function of P-gp in normal ageing human brains^60,61^. Since P-gp activity is ATP-dependent, our finding on its upregulation in both aged mouse and human brains may represent an adaptive response to impaired energy metabolism in ageing endothelial cells. In AD, a weakened adaptive P-gp expression upregulation may contribute alongside diminished transporter function in AD pathogenesis. For other AD-associated candidates we examined, some genes have known functional roles in the brain vasculature (e.g. Cd2ap^55^) and hence more direct interpretations regarding their differential expressions. The mechanistic linkage for the majority of other identified GWAS genes to neurodegeneration, however, remains unclear. It seems plausible that the expression changes of at least a subset of these genes may have pathogenic and/or adaptive roles in the aged brain vasculature yet to be uncovered.

Additionally, for a subset of aged mouse brain EC DEGs with important functional implications, we found similar changes in human AD brains compared to age-matched controls. Intriguing, these mostly had downregulation of expression. For instance, although upregulated in normal aged human brains, the reduced expression of GLUT1 glucose transporter gene Slc2a1 in aged mouse brains resembles that found in human AD^64–67^. GLUT1 knockdown in ECs has been shown to accelerate the development of pathology in the APP^Sw/0^ mouse model of AD^71^. We also found novel linkage of expression changes in aged brain and AD. Of note, endothelial downregulation of Mfsd2a is worth special highlight. MFSD2A serves as the major transporter for brain uptake of the essential omega-3-fatty acid docosahexaenoic acid across the BBB^72^. It is essential for BBB formation and maintenance, as mouse with MFSD2A knockout develop leaky BBB despite grossly normal vasculature morphology^73^. Of note, MFSD2A lipid transport function is required for inhibition of caveolae-mediated transcytosis across the endothelium^41^. The downregulation of MFSD2A may contribute to BBB breakdown and AD pathogenesis alongside other expression changes. Currently, AD pathogenesis and preclinical drug development rely heavily on mice harboring familial AD genes as experimental models and concerns often surround the generalizability of conclusions. Our finding that several key loss-of-function changes occur both in aged mouse and human AD brains supports the scientific ground of using aged mouse as an alternative model, with additional disease-mimicking factors depending on which research questions to address. For example, temporally controlled expression of mutant amyloid precursor protein and/or presenilin in aged mouse may be a more accurate model for AD therapeutics testing.

Ageing has long thought to be an irreversible process involving gene expression changes that impact diverse cellular processes. Strikingly, despite the complex subtype-dependent transcriptomic changes, we found a strong reversal effect of GLP-1R agonist treatment in all major EC subtypes that accompanies functional vascular benefit. This observation holds for DEGs of aged brain ECs associated with identified enriched functional pathways and AD. Although it is hard to dissociate the exact cellular pathways involved, our data suggest that correcting the expressions of immune/cytokine signaling, energy metabolism, and key BBB-regulatory genes may all play a role. Importantly, the finding also holds for the subsets of genes with concordant changes in human aged or AD brains (e.g. Slc2a1 and Mfsd2a). GLP-1R agonists have previously been found efficacious in several animal models of neurovascular and neurodegenerative diseases including stroke, AD, PD and ALS in protecting against neuronal death or degeneration^28–33^. Evidence also suggests that these findings may be generalizable to human patients. In a phase II PD trial, weekly subcutaneous injection of exenatide halted the deterioration of off-medication motor scores over a treatment period of 48 weeks, an effect that persisted even after a 12-week washout period^35^. Another phase II study in AD showed that six-month liraglutide treatment prevented the decline of brain glucose metabolism^34^. Given that ageing-associated NVU dysfunction represents a common component for AD and other neurodegenerative conditions, our study shows that reversal of ageing-associated changes in brain ECs and BBB protection is an important underlying mechanism that explains GLP-1R agonists' general applicability as potential disease-modifying therapeutics. Currently, further trials testing GLP-1R agonists are underway and may complement other approaches to the development of therapeutics for AD^74,75^.

The practical pharmacological approach we demonstrated complements the recent exciting discovery that young plasma infusion can partially reverse ageing-associated EC transcriptomic alterations^76^. In fact, apart from the reversal of expression changes in the aged brain endothelium, GLP-1R agonist treatment and young plasma infusion appear to share several functional benefits in animal models, such as amelioration of neuroinflammation^32,77^, rescuing synaptic plasticity deficits and cognitive benefits in aged or AD mouse^78,79^. Further studies are required to clarify the exact loci of actions for GLP-1R agonists. Due to multiple GLP-1R expression sites^32,36–38^ and capability of GLP-1R agonists to cross the BBB^80,81^, it remains to be determined whether the neurovascular benefits of GLP-1R agonist treatment may depend on vascular or glial cell GLP-1R, or acts by modulating immune cells or altering compositions in the peripheral circulation.

We wish to emphasize that the interpretation of transcriptomic data is often associative and not causal, and hence it is important to avoid false-positive discoveries or inflation of results. Despite the shallow sequencing depth and possibly limited by sample size, by adopting a relatively stringent criteria for defining significant DEGs, we sought to identify the most prominent functional pathways impacted and neurodegenerative disease-associations. It is also important to note that, despite the lack of significant overrepresentation, other neurovascular or neurodegenerative disease-associated genes with differential expression in the aged brain endothelium may still carry functional implications that are yet to be revealed (e.g. Nbeal1, associated with cerebral small vessel disease/stroke^54^ and most upregulated in aged arterial ECs). Moreover, in our study, whole-brain vasculature isolation was performed. For diseases with regional vulnerability (e.g. ALS, FTD, PD), we propose that targeted dissection of disease-affected regions for transcriptomic profiling of neurovascular and glial cells will be needed to uncover additional disease-associated expression changes in the aged brain.

## Methods

### Animal subjects

All experimental procedures were approved in advance by the Animal Research Ethical Committee of the Chinese University of Hong Kong (CUHK) and were carried out in accordance with the Guide for the Care and Use of Laboratory Animals. C57BL/6J mice were provided by the Laboratory Animal Service Center of CUHK and maintained at controlled temperature (22 – 23°C) with an alternating 12-hour light/dark cycle with free access to standard mouse diet and water. Male mice of two age groups (2 – 3 months old and 18 – 20 months old) were used for experiments. For the treatment groups, exenatide (5 nmol/kg/day, Byetta, AstraZeneca LP) was intraperitoneally (I.P.) administered starting at 17 – 18 months old for 4 – 5 weeks respectively prior to *in vivo* imaging and single-cell RNA sequencing experiments.

### Brain tissue dissociation and single cell isolation

We adapted a previously described dissociation protocols optimized for single brain vascular cells isolation from both young and old mouse brains^82^. Briefly, mice were deeply anesthetized and perfused transcardially with 20□ml of ice-cold phosphate buffered saline (PBS). Mice were then rapidly decapitated, and whole brains were immersed in ice-cold Dulbecco’s modified Eagle’s medium (DMEM, Thermo Fisher Scientific). The brain tissues were cut into small pieces and dissociated into single cells using a modified version of the Neural Tissue Dissociation kit (P) (130-092-628, Miltenyi Biotec). Myelin debris was removed using the Myelin Removal kit II (130-096-733, Miltenyi Biotec) according to the manufacturer’s manual. Cell clumps were removed by serial filtration through pre-wetted 70 μm (#352350, Falcon) and 40 μm (#352340, Falcon) nylon cell strainers. Centrifugation was performed at 300 xg for 5 minutes at 4°C. The final cell pellets were resuspended in 500 – 1000 μl FACS buffer (DMEM without phenol red (Thermo Fisher Scientific), supplemented with 2% fetal bovine serum (Thermo Fisher Scientific)).

### Single-cell library preparation, sequencing, and alignment

Single-cell RNA-seq libraries were generated using the Chromium Single Cell 3′ Reagent Kit v2 (10X Genomics, US). Briefly, single-cell suspension at a density of 500 – 1,000 cells/μL in FACS buffer was added to real-time polymerase chain reaction (RT-PCR) master mix aiming for sampling of 5000 – 8000 cells, and then loaded together with Single Cell 3′ gel beads and partitioning oil into a Single Cell 3′ Chip according to the manufacturer’s instructions. RNA transcripts from single cells were uniquely barcoded and reverse-transcribed within droplets. cDNA molecules were preamplified and pooled followed by library construction according to the manufacturer’s instructions. All libraries were quantified by Qubit and RT-PCR on a LightCycler 96 System (Roche Life Science, Germany). The size profiles of the pre-amplified cDNA and sequencing libraries were examined by the Agilent High Sensitivity D5000 and High Sensitivity D1000 ScreenTape Systems (Agilent, US), respectively. All single-cell libraries were sequenced with a customized paired end with single indexing (26/8/98-bp for v2 libraries) format according to the recommendation by 10X Genomics. All single-cell libraries were sequenced on a HiSeq 1500 system (Illumina, US) or a NextSeq 500 system (Illumina, US) using the HiSeq Rapid SBS v2 Kit (Illumina, US) or the NextSeq 500 High Output v2 Kit (Illumina, US), respectively. An average of 21017.2 mean reads per cell (ranging: 12665 to 51665) was obtained, which detected an average of 801.4 genes per cell (range: 540 to 1274). The library sequencing saturation was on average 77.89%. The data were aligned in Cell Ranger (v3.0.0, 10X Genomics, US).

### Single-cell transcriptomic data analysis

#### 1. Quality control and batch effect removal

Data processing and visualization were performed using the Seurat package (v3.0.1)^83^ and custom scripts in R (v3.5.1). The raw count matrix was generated by default parameters (with the mm10 reference genome). There were 63300 cells in the primary count matrix. Genes expressed by fewer than 3 cells were removed, leaving 19746 genes in total. Among these genes, 4000 high-variance genes were identified by the Seurat *FindVariableFeatures* function. To remove batch effects between different sequencing batches, we used a canonical correlation analysis (CCA)-based integration function in Seurat. The high-variance genes were used to find anchor genes as a stable reference to eliminate batch effect from the first 30 dimensions from CCA. After batch effect removal, the reassembled dataset was filtered to exclude low quality cells by the following criteria: 1. Lower than 5% or higher than 95% UMI count or gene count. 2. The proportion of mitochondrial genes > 20%. For dimensionality reduction, principal component analysis (PCA) was applied to compute the first 60 top principal components. Clustering was carried out by the Seurat functions *FindNeighbors* and *FindClusters*. The *FindNerighbors* function constructed a shared nearest neighbor (SNN) graph based on the first 50 principal components. Modularity optimization was then performed on the SNN results for clustering (resolution parameter: 1.6). We employed t-distributed stochastic neighbor embedding (t-SNE) to visualize the clustering results and included 52 initial clusters for further analysis after removal of 3 clusters with abnormally high mitochondrial gene percentages.

#### 2. Primary cell type and endothelial cell subtype identification

To identify primary cell types, we employed known cell type-specific marker genes and examined their expression levels among all 49 initial clusters included. We further excluded clusters with a dual-high expression of two or more cell type-specific marker genes. These included a cluster with high expression of both endothelial cell and pericyte markers, corresponding to contamination of pericytes by endothelial cell fragments also reported in previous literature^1,84,85^. The remaining clusters were classified into 16 primary cell types (see Fig. 1 and Supplementary Fig. 1). To allow identification of EC subtypes, we employed a probabilistic method, *CellAssign*^39^, to classify each EC into one of the six principal subtypes using 267 variable genes along the arteriovenous axis from previously published literature as prior knowledge^1^. Each cell was assigned to the subtype with the highest likelihood (see Supplementary Table 1 for full lists of the 267 genes), and the algorithm was run 3 times independently. Only endothelial cells with identical subtype assignment for all 3 runs were included for further analysis (11283 out of 12357, 91.3%).

#### 3. Differentially expressed gene calculation

After batch effect removal, quality control and cell type/subtype classification, gene count normalization and high-variance gene identification were applied to the raw data of 43922 cells retained for further analysis. The Seurat *FindMarkers* function and the MAST package (v1.8.2) were employed for calculation of differentially expressed genes (DEG) with associated raw *P*-value, false discovery rate (FDR)-adjusted *P*-value and magnitude of change expressed in natural log of fold change (*lnFC*) for each cell type/subtype. We define significant DEGs as those fulfilling FDR-adjusted *P*-value < 0.05 and an absolute value of *lnFC* exceeding 0.1.

#### 4. Pathway enrichment and GWAS-disease gene overrepresentation analysis

We used GeneAnalytics, an online universal gene functional analysis tool, which contained more than a hundred data sources for pathway enrichment analysis. Significant DEGs were converted to human gene orthologs semi-automatically. Pathways with prominent functional implications, including blood-brain barrier (BBB)-related pathways, immune/cytokine signalling pathways, respiratory electron transport chain / ATP synthesis and glucose/energy metabolism pathways were identified manually from literature review and summarized. We merged enrichment results on TGF-β and VEGF signaling from different sources (i.e. GeneCards, Wikipathways) and presented only the most significant item in the main text/figure.

For neurovascular or neurodegenerative diseases examined (Supplementary Table 2), we compiled GWAS-identified genes associated with disease risk or phenotypic traits from the GWAS catalog and selected recent GWAS studies (see Supplementary Table 2 for full lists of diseases, associated genes and sources). We then found the intersection of the human orthologs of significant DEGs for each cell type / subtype-disease combination and performed hypergeometric test for overrepresentation by the *phyper* function in R.

#### 5. Analysis of GLP-1R agonist treatment group scRNA-seq data

Additional single-cell RNA-sequencing data from the GLP-1R agonist treatment group were processed in parallel with young adult and aged mouse groups by the same pipeline. From the treatment group, we obtained 7239 brain ECs in total, with the following subtype distribution: aEC1, *n* = 978; aEC2, *n* = 1155; capEC, *n* = 1728; vcapEC, *n* = 1851; vEC, *n* = 1043; avEC, *n* = 484. Single-proportion *z*-test was employed to test whether proportions of genes with reversed expression changes due to treatment was higher than chance level. The proportions of upregulated and downregulated genes were calculated in aged versus young group (P_ovy_up_ and P_ovy_down_, only for DEGs with FDR-adjusted *P*-value < 0.05), and aged exenatide-treated versus untreated group (P_evo_up_ and P_evo_down_, for all genes). Background chance of reversal of expression changes of both up- and downregulated DEGs was then calculated as P_ovy_up_ × P_evo_down_ + P_ovy_down_ × P_evo_up_ (i.e. assuming independence of probability of up- or downregulation by treatment in relation to ageing-associated changes). Background chance of downregulation of ageing-upregulated DEGs was simply P_evo_down_, while upregulation of ageing-downregulated DEGs was P_evo_up_. One-sided, one-proportion *z*-test was used to assay whether proportions of genes with reversal of expression changes was higher than chance level. Linear regression was performed and the slope of best linear fit, *r*^*2*^ and *P*-value for the regression coefficient were obtained. Selection of BBB-regulatory, immune/cytokine signalling, energy metabolism pathway-associated genes were based on annotations from pathway enrichment analysis by GeneAnalytics.

### Human brain bulk RNA sequencing dataset analysis

Human brain bulk RNA sequencing data (de-identified) from different ages were obtained from the Genotype-Tissue Expression (GTEx) portal, excluding data from the cerebellum and cervical spinal cord. 362 samples from 61 subjects were divided into two age groups (< 60 and ≥ 60 years old). Comparisons of expression levels (expressed in TPM) for all genes were carried out across the two age groups by *t*-test with FDR-adjustment for multiple comparisons. For the Allen Brain Institute Aging, Dementia and Traumatic Brain Injury study dataset (Adult Changes in Thought (ACT) study cohort)^62,63^ of human brain bulk RNA sequencing from the Allen Brain Institute, 111 samples from 30 patients were defined as controls with the following criteria: 1. Annotated as “no dementia” based on the DSM IV diagnostic criteria. 2. No history of traumatic brain injury. 58 samples from 16 patients were classified as human AD brain samples with the following criteria: 1. Diagnosed with Alzheimer's disease dementia based on the DSM IV diagnostic criteria. 2. No history of traumatic brain injury. All AD subjects included had pathologically confirmed AD with Braak Staging: I, *n* = 2; II, *n* = 2; III, *n* = 1; IV, *n* = 2; V, *n* = 5; VI, *n* = 4. Comparisons of expression levels (expressed in RPKM) for all genes were carried out across the two groups by *t*-test with FDR-adjustment for multiple comparisons. EC-enriched genes were defined as genes with *lnFC* > 0.7 (i.e. at least two-fold higher) and FDR-adjusted *P*-value < 0.05 in ECs relative to all other cell types from the mouse brain scRNA-seq data. Expression changes for the human orthologs of all EC-enriched genes in the GTEx and ACT datasets were calculated by unpaired *t*-test with FDR-adjustment and directionality of changes were compared to corresponding aged mouse brain DEGs. One-sided, two-proportion *z*-test was used to assay if among the human orthologs of significant DEGs of aged mouse brain EC subtypes, the proportion with significant changes between the two human brain age groups was higher than that among all background genes.

### Cranial window implantation

To permit *in vivo* two-photon microscopic imaging of the brain, animals underwent a cranial window implantation surgery. Prior to surgery, mice received one dose of dexamethasone (2 mg/kg, S.C.) to reduce inflammation. Animals were anesthetized with ketamine (100 mg/kg, I.P.) and xylazine (10 mg/kg, I.P.). Body temperature was maintained at 37□°C by a homeothermic heating pad system. During the surgery, mice were fixed in a stereotaxic instrument for rats and mice (New standard™ 51500D, Parkland Scientific). A cranial window (5 mm diameter) was made over the left somatosensory cortex. A customized chamber frame was placed around the opened skull and fixed with cyanoacrylate gel superglue (Loctite 454). The exposed cortex was covered with a 5-mm glass coverslip fixed by cyanoacrylate gel superglue. The rest of the cranial window margin and skull area were filled with dental cement. All procedures were performed carefully to prevent damage to the cortex and vessel structures. Following surgery, mice were given one dose of enrofloxacin (Baytril, 5 mg/kg S.C.) for prevention of infection and buprenorphine (Temgesic, 0.1 mg/kg S.C.) for analgesia.

### In vivo two-photon imaging assay of blood-brain barrier integrity

To assay BBB permeability, mice were anesthetized with intraperitoneal ketamine (100 mg/kg) and xylazine (10 mg/kg) and placed on a head-fixing apparatus under a custom-modified two-photon microscope (Scientifica, Uckfield, UK) with a Ti:Sapphire femtosecond laser (Mai Tai DeepSee, Spectra-Physics), galvo-resonant scanners and a Nikon ×16/0.80NA water-immersion objective. A dye mixture containing 12.5□mg/ml fluorescein isothiocyanate (FITC)-conjugated dextran (70 kDa molecular weight, Sigma-Aldrich) and 6.25□mg/ml tetramethylrhodamine (TRITC)-conjugated dextran (40 kDa molecular weight, Thermo Fisher Scientific) dissolved in PBS was administered (4 ml/kg body weight, i.v.) via tail vein injection. *In vivo* image stacks were then acquired 15 – 35 minutes post-injection, with 930 nm excitation wavelength, while the emitted fluorescent signals were detected by photomultiplier tubes through a 520/20 band-pass filter for FITC-conjugated dextran and a 609/34 band-pass filter cube for TRITC-conjugated dextran. Laser power under the objective was kept under 50 mW for all imaging sessions. Three image stacks for each mouse were acquired in custom-modified ScanImage (v5.3.1, Vidrio Technologies, US) in MATLAB R2017b (Mathworks, US) and used for the quantification of TRITC-conjugated dextran extravasation. Each image stack had a field of view of 417 μm × 417 μm and consisted of 300 imaging planes evenly distributed over 239 μm, starting from 50 μm below the cortical surface. 6 images were acquired and averaged for each plane. Measurement of fluorescent signals and volumetric reconstruction of extravasated TRITC-conjugated dextran was carried out using the Imaris software (v6.4, Bitplane, Belfast, UK). To separate intra- and extravascular spaces, volumetric reconstruction of the vasculature was performed using the FITC-conjugated dextran images (i.e. large molecular weight 70 kDa dextran which remained in vessels). Volumetric reconstruction of the TRITC-conjugated dextran in extravascular spaces was then performed and quantified. Volume rendering was performed in normal shading mode. Artefacts were removed by applying gaussian filter. During 3D reconstruction, surfaces of vessels were generated by background subtraction with maximum surface detail limited to 0.814 μm. Differences between experimental groups were compared by one-way ANOVA followed by Tukey’s multiple comparison test in the Prism software (v6, GraphPad, California, US).

## Data Availability

The raw and processed RNA sequencing data from this study will be deposited in the NCBI Gene Expression Omnibus database upon acceptance of the manuscript. R codes for sequencing data analysis are available upon reasonable request to the corresponding authors.

## Acknowledgments

We thank members of the Ko laboratory for helpful discussions; Rossa Chiu and Dennis Lo for generous support and access to sequencing facilities, David Attwell and William Wu for insightful advice; Becky Yung, Anki Miu, Rebecca Chau, Pauline Kwan and Rachel Hui for providing administrative support to the project. This work was in part funded by the Faculty Innovation Award (FIA2017/B/01), the Gerald Choa Neuroscience Center, Margaret K. L. Cheung Research Centre for Parkinsonism Management and the Chow Yuk Ho Technology Center for Innovative Medicine of the Faculty of Medicine, the Chinese University of Hong Kong (CUHK); the Area of Excellence Scheme of the University Grants Committee (AoE/M-604/16).

## Author Contributions

L.Z., Z.L. and J.S.L.V. carried out scRNA-seq experiments. Z.L. analyzed the scRNA-seq data with advice from H.K.. L.Z. performed *in vivo* two-photon imaging with assist by Z.L. and S.K.H.S.. L.Z. analyzed the *in vivo* imaging data with input from L.Y.C.Y.. Z.L. analyzed human brain transcriptome data with input from W.L.N. and H.K. H.M.L. reviewed human brain expression databases. H.M.L. and J.H. contributed to study design and data interpretation. H.Y.E.C., H.C.S., Y.T. and W.J.L. contributed to technical discussions and data interpretation. X.T. and Y.H. contributed to *in vivo* experimental protocol establishment. Z.L. and X.C. prepared manuscript figures. Z.L., H.M.L., V.C.T.M. and H.K. wrote the manuscript with input from all authors.

---

**Supplementary Fig. 1.**
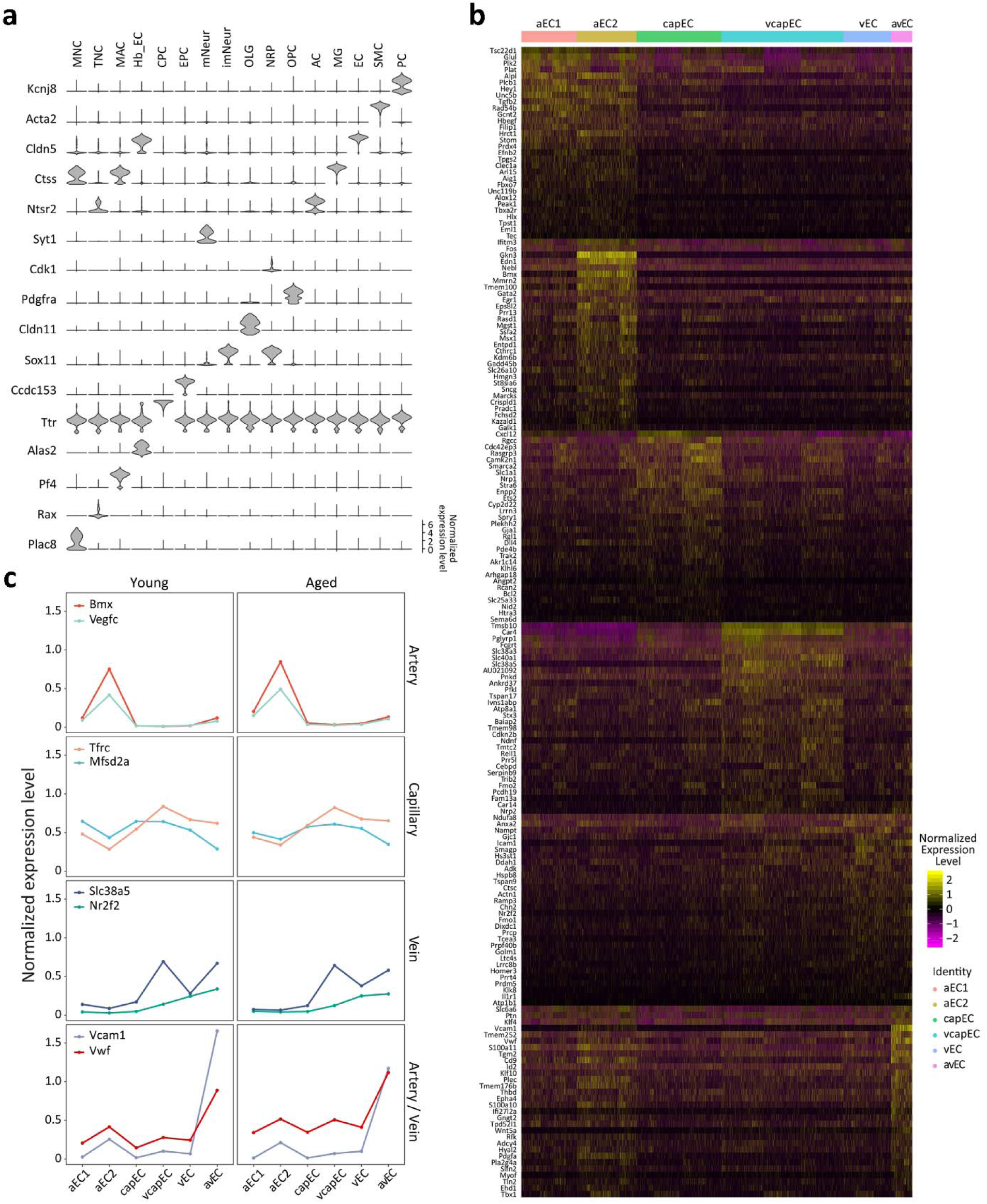
**a**, Violin plots of the expression patterns of marker genes for major cell type clusters identified. Each column corresponds to one primary cell type. Cell type abbreviations: same as **Fig. 1b,** Heatmap showing the expression patterns of variable genes along the arteriovenous axis employed for classification of EC subtypes (see Methods). Up to 30 genes are shown for each EC subtype. **c,** Relative average expression levels of marker genes of artery/arteriole (Bmx and Vegfc), capillary (Tfrc and Mfsd2a), vein (Slc38a5 and Nr2f2), and artery/vein (Vcam1 and Vwf) for the EC subtypes.

**Supplementary Fig. 2.**
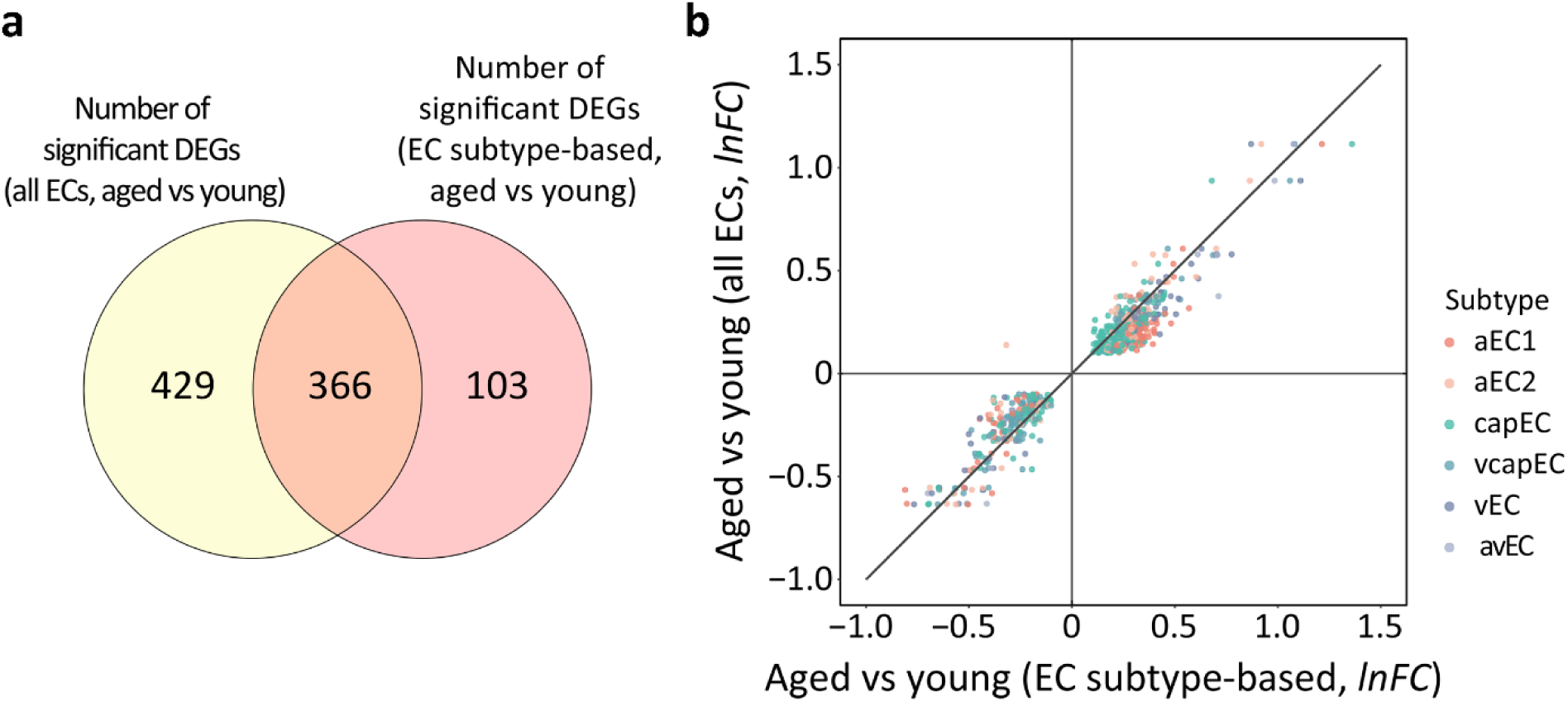
**a,** Venn diagram of EC subtype DEGs and pooled EC DEGs. The expression changes of a substantial proportion (22.0%, 103 out of 469) of DEGs would not be detected without subtype classification. **b,** *lnFC* in EC subtypes (x-axis) plotted against *lnFC* in all ECs pooled (y-axis) for DEGs significant in both subtype- and pooled EC-based comparisons across age groups. Note that genes with significant differential expression in more than one EC subtype have multiple points. For up- or downregulated DEGs, the majority of points (60.1%, 424 out of 706, *P* < 1 × 10^−4^) fall below or above the line with unit slope respectively.

**Supplementary Fig. 3.**
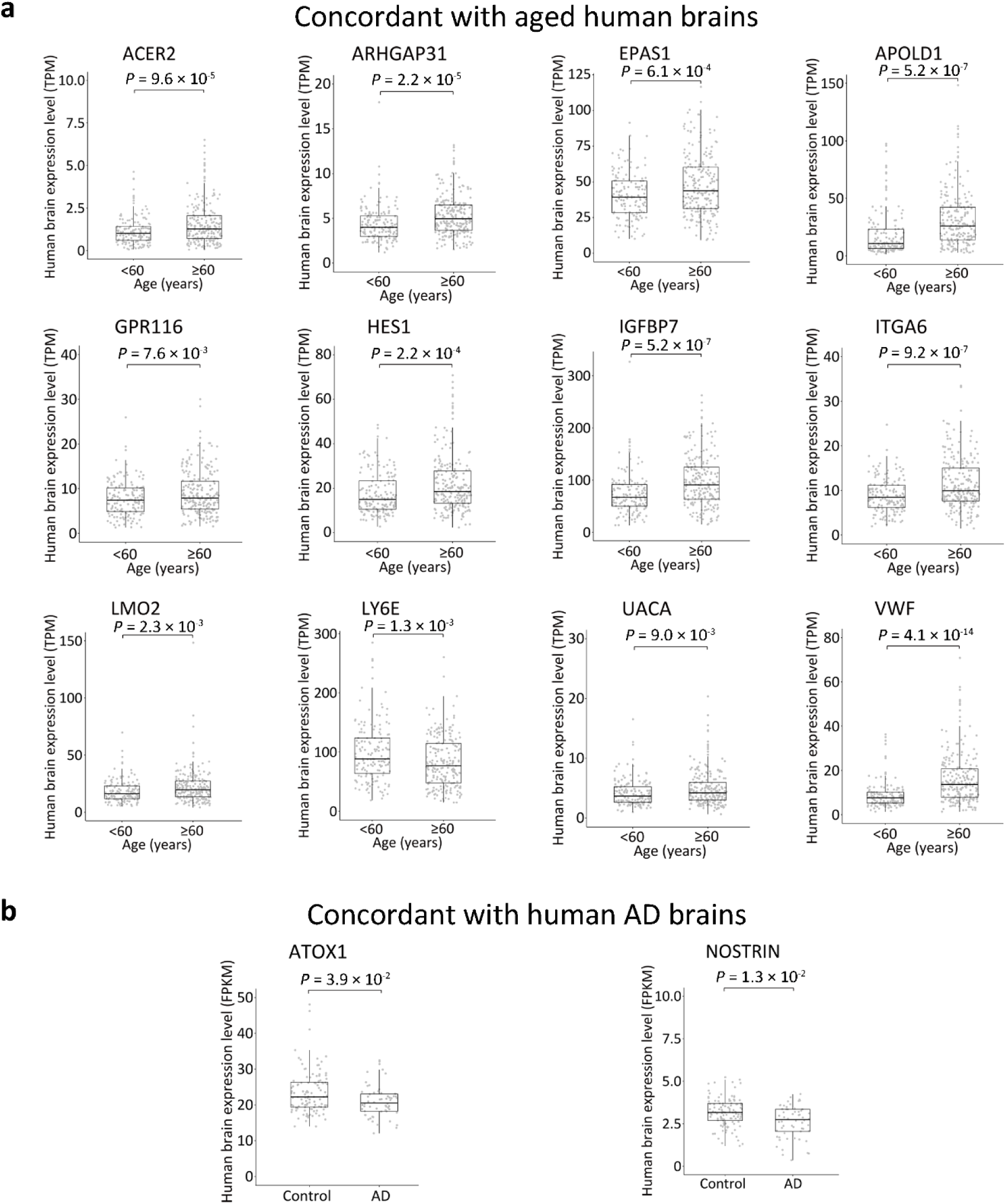
Additional boxplots for the human orthologs of aged mouse brain EC DEGs with concordant expression changes in human aged (**a**) or AD brains (**b**).

**Supplementary Fig. 4.**
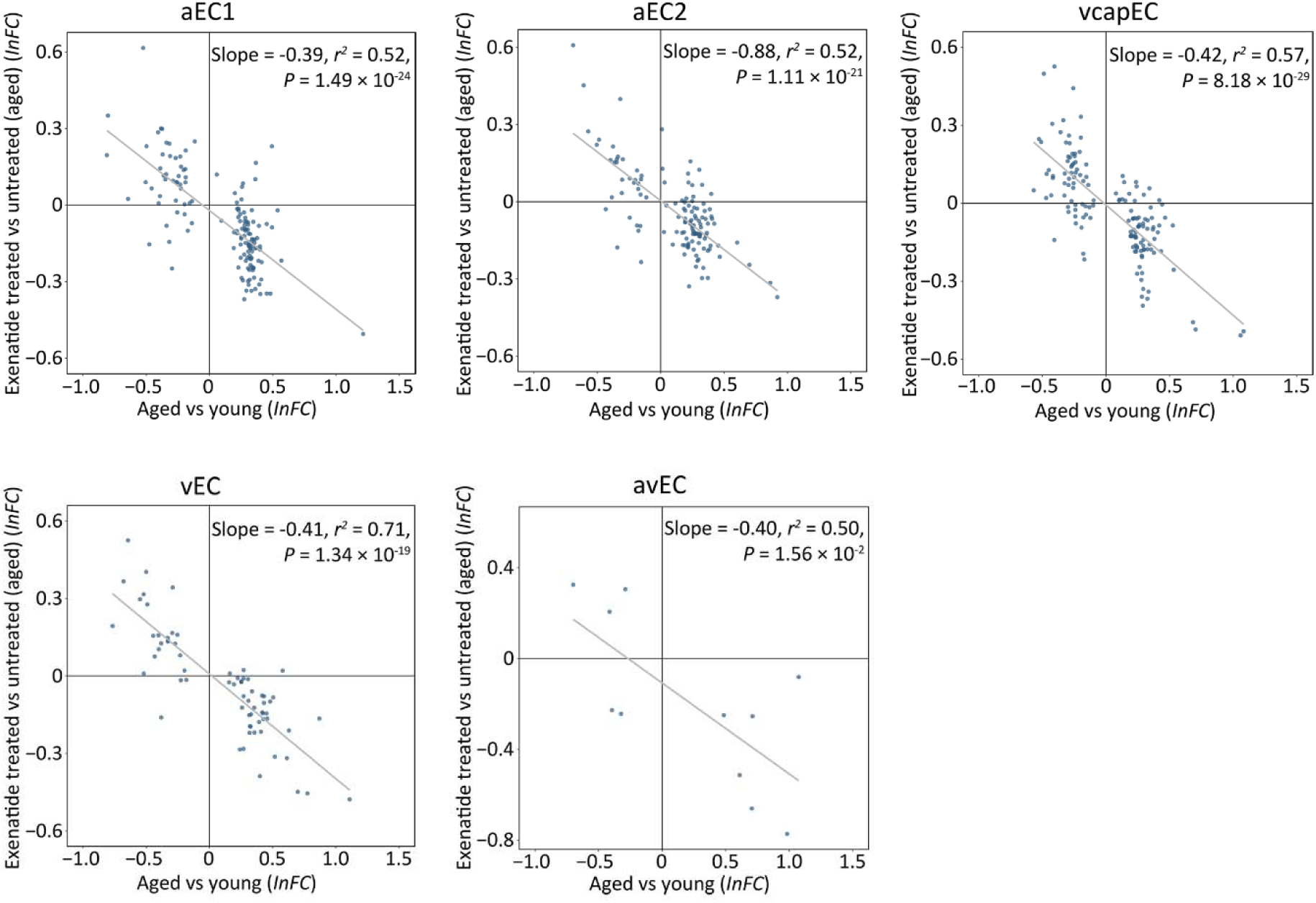
Reversal of ageing-associated transcriptomic changes by exenatide treatment in the EC subtypes.

