## Supplementary Table 1 for "Zonation-dependent single-endothelial cell transcriptomic changes in the aged brain"

| EC subtype | Marker genes (source: Nature. 2018 Feb 22;554(7693):475-480.) |
| --- | --- |
| aEC1 | Sncaip/Alpl/Alox12/B4galnt1/Glul/Syt15/Tpgs2/Fam198b/Antxr1/Eml1/Gm609/Rnf144a/Stom/Prdx4/Aig1/Arhgef25/Stc1/Plat/Peak1/Tec/Efnb2/Fbxo7/Unc5b/Tpst1/Hey1/Cpm/Hrct1/Unc119b/Hbegf/Tbxa2r/Spry4/Scube2/Plcb1/Hlx/Arl15/Tsc22d1/Filip1/A430090L17Rik/Plk2/Slc36a1/Sez6/Rad54b/Gcnt2/Clec1a/Tgfb2/Slc12a5 |
| aEC2 | Egr1/Mgst1/St8sia2/Entpd1/Zbp1/Cables2/Cthrc1/Msx1/St8sia6/Cbr3/Irf6/Lcat/Olfml2a/Slc27a3/Col18a1/Fchsd2/Prkcd/Ifitm3/Mmrn2/Gata2/Fos/Marcks/Edn1/Kdm6b/Eps8l2/P2ry2/Pfkfb3/Snx10/Ssfa2/Crispld1/Prr13/Atl2/Galk1/Kazald1/Dsp/Sncg/Rasd1/Nebl/Gadd45b/Lama3/Bmx/Gkn3/Tmem100/Ssbp2/Hmgn3/Slc26a10/Pradc1 |
| capEC | Dll4/Rasgrp3/Akr1c14/Slc25a33/Smarca2/Ets2/Rcan2/Sema6d/Gja1/Enpp2/Angpt2/Col4a3/Prdm1/Cdc42ep3/Hdac9/Pde4b/Arhgap18/Bcl2/Htra3/Gpr85/Cyp2d22/Itga4/Trak2/Nrp1/Camk2n1/Nid2/Lrrn3/Plekhh2/Rgcc/Stra6/Rgl1/Pcx/Slc1a1/Osgin1/Klhl6/Cxcl12/Spry1 |
| vcapEC | Nrp2/Frmd5/Tmem98/Pnkd/Atp8a1/Tmem37/Stx3/Pak1/Pitpnm2/Car14/Jak3/Tspan17/Serpinb9/Unc13c/Fcgrt/Pglyrp1/Kcp/Slc40a1/Slc38a3/Ctla2b/Tmtc2/Irf5/Nos2/Rell1/Hcn2/Coro2b/Baiap2/Rab4a/Pfkl/Trib2/Prr5l/Tmsb10/Ndnf/Car4/Slc38a5/Fmo2/Cebpd/St6gal1/AU021092/Pcdh19/Igsf5/Odc1/Fam13a/Tbx4/Ankrd37/Pmaip1/Grb7/Ivns1abp/Cdkn2b |
| vEC | Dixdc1/Nr2f2/Lbp/Adk/Ltc4s/Tspan9/Tcea3/Ndufa8/Lrrc8b/Chn2/Prcp/Icam1/Il1r1/Tbc1d2b/Nckap5/Gm7694/Bst1/Sdk1/Lcn2/Fmo1/Ctsc/Gpr182/Gm5127/Cysltr1/Hspb8/Prpf40b/Rp1/Mxra8/Prdm5/Itga3/Hs3st1/Anxa2/Golm1/Homer3/Atp1b1/Nampt/Prrt4/Sytl2/Gjc1/Actn1/Klk8/Ddah1/Smagp/Ramp3 |
| avEC | Tmem176b/Tgtp1/Id2/Ptn/Gngt2/Tpd52l1/Frem2/Hyal2/Ifi27l2a/Tln2/Slc6a6/Car7/Vcam1/Klf4/Rfk/P2ry1/S100a10/Ehd1/Plec/S100a11/Pla2g4a/Cd9/Pdgfa/Ntn1/Thbd/C530008M17Rik/Aldh1a3/Kcnb1/Tnfrsf11a/Klf10/Carhsp1/Aldh1a1/Wnt5a/Icosl/Cfb/Myof/Adh1/Epha4/Tgm2/Slfn2/Vwf/Adcy4/Tmem252/Tbx1 |
