## Supplementary Table 2 for "Zonation-dependent single-endothelial cell transcriptomic changes in the aged brain"

| Disease / condition | GWAS genes | Source(s) of GWAS genes |
|---|---|---|
| White matter hyperintensity burden / cerebral small vessel disease | TRIM65/EFEMP1/TRIM47/PLEKHG1/PDCD11/NEURL1/SH3PXD2A/PMF1/NBEAL1/NRCAM/DAB1/PTPRD/EVL/ZNF16/CHRM3/RASSF3/COL4A2/AHCYL2/C10orf143/PCDH7/QRICH2/KCNB1/DPPA3/ARID1A/VCAN/ITPRID1/PMF1-BGLAP/C1QL1/GAS1 | GWAS Catalog; Circ Cardiovasc Genet. 2015 Apr;8(2):398-409. |
| Stroke | UBE2E3/FOXF2/OPRM1/MPDZ/CFL2/SYNE2/ALCAM/CDH6/CASC10/NINJ2/SPRY2/F5/PITX2/F2/LRAT/SH2B3/HDAC9/FUNDC2/ALDH2/PMF1BGLAP/ABCC1/ZFHX3/FGB/LRCH1/PATJ/CHD3/SLC44A2/CASZ1/PROCR/FAF1/CDK6/SLCO1B1/ILF3/CDKN2C/PRPF8/PDE3A/ADAMTS13/SMARCA4/DAB1/AGBL1/KIF26B/COX7A2L/SWAP70/CDKN1A/PTPRD/TSPAN15/ALDH1A2/IRX1/SLC26A11/PTPRG/KCNK1/HPS4/MYRIP/C10orf143/PCDH7/QRICH2/CDC5L/CRISP2/KCNK3/DPPA3/ARID1A/NRCAM/LAMA1/WNT2B/TTBK1/RPH3A/ONECUT2/LMNA/RAB19/VWF/FUT8/ADGRF1/KAT2B/FGG/PITX2/ATXN2/TWIST1/PMF1/FGA/ABO/SSPN/TCF7L2/ANK2/FURIN/SH3PXD2A/RHAG/CDH13/ATP6V1G3/KNG1/SLC22A7/APOD/C17orf102/USP38/TNFRSF21/RASEF | GWAS Catalog; Lancet Neurol. 2016 Jun;15(7):695-707. |
| Parkinson's disease | GBA/NUCKS1/SLC41A1/ITPKB/SIPA1L2/IL1R2/TMEM163/CCNT2/SCN3A/STK39/SATB1/NCKIPSD/CCDC71/ALAS1/TLR9/DNAH1/BAP1/PHF7/NISCH/STAB1/ITIH3/ITIH4/MCCC1/TMEM175/DGKQ/FAM200B/CD38/FAM47E/SNCA/ANK2/CAMK2D/ELOVL7/ZNF184/HLAQA1/KLHL7/NUPL2/GPNMB/CTSB/MICU3/SORBS3/PDLIM2/C8orf58/BIN3/SH3GL2/FAM171A1/BAG3/DLG2/LRRK2/OGFOD2/GCH1/TMEM229B/GALC/VPS13C/COQ7/ZNF646/KAT8/TOX3/ATP6V0A1/PSMC3IP/TUBG2/ARHGAP27/CRHR1/SPPL2C/MAPT/STH/KANSL1/SYT4/LSM7/DDRGK1/CHMP2B/KRTCAP2/CNTN1/RAB29/CCDC62/BST1/INPP5F/WNT3/HLADRB1/RIT2/GAK/NDUFAF2/NSF/DCUN1D1/HLADQB1/BCKDK/LAMTOR2/SLC2A13/LHFPL2/TIAL1/LZTS3/MAP4K4/MMRN1/TPM1/GPR65/SYT17/HLADRA/IP6K2/PLPPR1/ITGA8/TBC1D5/PMVK/SCN2A/ITIH1/KLHDC1/SLC50A1/COL3A1/CA8/PLEKHM1/CCN6/SREBF1/TCEANC2/AGAP1/CYP17A1/CCDC82/TMC3/PAM/ZNF165/GBF1/KTN1/MX2/CAB39L/TAS1R2/QSER1/COL13A1/TMPRSS9/ZP3/CNKSR3/TRAPPC2L/COL5A2/MDGA2/BRINP1/SPTSSB/KCNIP4/STAP1/TRPS1/UNC13B/ANO5/RBMS3/ITGA2B/CLRN3/DSG3/ODAPH/MARCH3/PRDM15/ISM1/LTK/SEMA5A/ATF6/SP1/CTC1/AAK1/SYT10/FAM47ESTBD1/FAM126A/RAB25/KCNN3/WBP1L/MED13/PRSS53/PAX7/PRRG4/PABPN1L/WNT9A/IGSF11/FDFT1/HTR2A/GFPT2/OCA2/CHL1/LMNB1 | GWAS Catalog; Nat Genet. 2017 Oct;49(10):1511-1516. |
| Corticobasal degeneration | MAPT /MOBP/SOS1/LINC02210-CRHR1/SPPL2C/KIF13B/DUSP4/TSPEAR | GWAS Catalog; Nat Commun. 2015 Jun 16;6:7247. |
| Progressive supranuclear palsy | CD8B/IRF4/IL2/IL21/STX6/EIF2AK3/MOBP/MAPT/BMS1/SLCO1A2/EXOC2/KANSL1/NSF/WNT3/TRIM11/RUNX2/SP1/ASAP1/PIK3C2G | GWAS Catalog; Nat Genet. 2011 Jun 19;43(7):699-705. |
| Multiple systems atrophy | FBXO47/ELOVL7/EDN1/CRHR1/MAPT/RREB1/ASB1/WNT3/MDGA2/CDH4/LOXL4/EBF2/CARD6/USP31/LMX1B/FBN2/XDH/SLC28A3/FOXL1/MPP6/ANKFN1/FOXN3/ENY2/NDE1/ARID1B/ASCC3/RIMS2/TENM2/DDX18/ASXL3/RASGRP3/ARHGAP44/ACOT11/ARL17A/RPL19/PLEKHM1/KANSL1/LINC02210CRHR1/PKHD1L1/PYROXD2/HS3ST2/THSD7B | GWAS Catalog; Neurology. 2016 Oct 11; 87(15): 1591–1598. |
| Frontotemporal dementia | RAB38/CTSC/HLADRA/BTNL2/TMEM106B/VWDE/GFRA2/CEP131/WASHC5/NDUFAF8/TEPSIN/COL28A1/ST18/ATP9B/TNIK/RRBP1/BANF2/NDUFS1/C8orf49/NEIL2/FUT10/CFAP54/SEM1/DLD/HNF1B/CHL1/ZCCHC24/ACTN3/PALM2/SFRP4/STARD3NL/DMRT2/SLC38A10/UNC13A/TRMT11/ZFHX3/NECTIN2/LRRC37B/PRPF40A/SHANK3/SP1/FAT4 | GWAS Catalog; Lancet Neurol. 2014 Jul;13(7):686-99. |
| Amyotrophic lateral sclerosis | RPSA/SLC25A38/C9orf72/MOBP/SCFD1/POLDIP2/TMEM199/SARM1/KCNN1/KIF5A/MOB3B/UNC13A/ADGRD1/SUSD2/IDE/TBK1/C5orf30/OPCML/KIFAP3/TIAM1/ITGA9/CTNND2/ZNF746/TNIP1/ALCAM/CAMTA1/DPP6/WASHC5/ETNPPL/OLFM4/METTL21A/MASP1/CENPV/ATXN3/ANK3/PSD3/CPNE4/NPEPPS/KIAA0513/ARAP2/ABCG1/STK36/ZFYVE26/PBDC1/SLC9A9/NPS/LAMA3/CLVS1/KCNMB2/INPP4B/EFEMP | GWAS Catalog; Nat Genet. 2016 Sep;48(9):1043-8. Neuron. 2018 Mar 21;97(6):1268-1283.e6. |

| | | |
|---|---|---|
| | 1/NME9/ARHGEF2/SLC25A12/ERBB4/HOXD10/KALRN/KCNS3/IFRD1/SYNPO2/TBXAS1/ADAMTSL1/TMEM132B/NOG/CREB5/ITPR2/FOLH1B/TSPAN9/TFAP2A/STON1/PTH2R/HADH/LAMA2/GRID1/NEDD4L/MRAS/TBC1D1/ATXN1/SELL/TRPM8/CALN1/ASIC2/PROCR/ZFP64/SQLE/PTPRF/CNOT2/C3orf56/ANXA3/RGS6/NFATC2/SUSD1/NRXN3/EPB41/CCSER1/PDLIM5/TAF8/PDGFRL/RBM19/CHODL/CNTN4/SCN7A/CTDSP1/PCSK5/ATP2B2/GSE1/DISC1/NT5C1A/SLC39A11/MORN2/COMMD10/CCDC192/ANKS1B/NFASC/MACROD2/ANKRD29/FHDC1/MIR99AHG/LDHC/DACH1/ABCC12/PPP2R2D/CFAP410/ABCC12/ST3GAL3/OSTC/PIGL/MAP3K7/DOCK1/MYOM2/PNPT1/ZBTB40/C17orf67/SEC16B/EGR1/SLC18A1/CALML3/STON1GTF2A1L/LEF1/WAPL/C1orf112/SPP2/KDM4A/LRRC8C/PLXNA1/XIRP2/VIL1/LIPC/RBMS1/ALDH1A2 | |
| Huntington's disease | FAN1/SOSTDC1/ISPD/ADGB/PTPRM/OR10A2/KIF9/BRF1/ADD1/TENM2/PTDSS1/EFR3A/MSH3/TRPM1/RNASET2/PRDM9/MTMR10/ATRNL1/GFRA1/ST8SIA6 | GWAS Catalog; Lancet Neurol. 2017 Sep;16(9):701-711. |
| Alzheimer's disease | PPOX/B4GALT3/ADAMTS4/NDUFS2/FCER1G/APOA2/CR1/CR1L/BIN1/ERCC3/GPR17/INPP5D/HESX1/MICB/C4A/C4B/PRRT1/EGFL8/AGPAT1/RNF5/AGER/PBX2/GPSM3/NOTCH4/TSBP1/BTNL2/HLA-DRA/HLA-DRB5/HLA-DRB1/HLA-DQA1/HLA-DQB1/HLA-DQA2/HLA-DQB2/HLA-DOB/TAP2/PSMB8/PSMB9/VPS52/TREML1/TREM2/CD2AP/ADGRF2/TRIM4/AZGP1/ZKSCAN1/ZSCAN21/COPS6/MCM7/AP4M1/TAF6/CNPY4/MBLAC1/LAMTOR4/C7orf43/GAL3ST4/GPC2/STAG3/CASTOR3/PVRIG/PILRB/PILRA/ZCWPW1/MEPCE/PPP1R35/C7orf61/TSC22D4/NYAP1/AGFG2/LRCH4/SAP25/FBXO24/MOSPD3/TFR2/GIGYF1/EPHB4/SLC12A9/FAM131B/ZYX/EPHA1/TAS2R60/ARHGEF35/OR2A7/CNTNAP2/TRIM35/PTK2B/CHRNA2/EPHX2/CLU/SCARA3/CCDC25/TCN1/MS4A3/MS4A2/MS4A6A/MS4A4E/MS4A4A/MS4A6E/PICALM/HIKESHI/SORL1/SLC24A4/RIN3/ADAM10/MINDY2/RNF111/SLTM/TPM1/LACTB/RAB8B/APH1B/CCDC189/RNF40/HSD3B7/STX1B/STX4/ZNF668/ZNF646/PRSS53/BCKDK/KAT8/PRSS8/PRSS36/ITGAX/MINK1/CHRNE/C17orf107/RNF167/CAMTA2/INCA1/KIF1C/ZFP3/ZNF232/USP6/SCIMP/RABEP1/NUP88/GNGT2/PHOSPHO1/ZNF652/ALPK2/FGF22/WDR18/GRIN3B/CNN2/ABCA7/ARHGAP45/POLR2E/GPX4/ZNF223/ZNF225/IGSF23/PVR/CEACAM19/CEACAM16/BCL3/CBLC/BCAM/NECTIN2/TOMM40/APOE/APOC1/APOC4/APOC2/CLPTM1/RELB/CLASRP/ZNF296/GEMIN7/MARK4/PPP1R37/NKPD1/TRAPPC6A/BLOC1S3/EXOC3L2/CKM/KLC3/ERCC2/EML2/FBXO46/DMPK/DMWD/IRF2BP1/CD33/SIGLECL1/VSIG10L/CSTF1/CASS4/RTF2/FAM209B/NECTIN2/APOC1P1/IL6R/MMP3/MMP12/ACKR2/KRBOX1/CYP8B1/UNC5CL/CCRL2/OARD1/CEACAM20/HNF4G/SUCLG2/BCAS3/BZW2/SPON1/FRMD4A/MTHFD1L/CR1/RBFOX1/ABI3/PLCG2/CLU/ATP5F1C/TMC5/CACNA1G/SYNJ1/GLIS3/CST1/GMNC/GPR141/ARL17B/PFDN1/NKAIN3/SDR42E2/FERMT2/PHF14/CTNNA2/ANKRD55/SAP30L/CELF1/NFIC/ITSN2/KDM1B/TMEM132C/MYO16/CLMN/MRPL58/IQCK/TGM6/ZNF292/CYYR1/EDAR/KAT8/PTPRG/LARS/TNRC6A/RAPSN/DSG2/BHMG1/OSBPL6/F13A1/FBXL7/CRADD/ACE/CEACAM19/ADGRF2/CASTOR3/SPI1/ZNF232/SNX9/COL12A1/CCDC89/FMN2/GPC6/VSTM2A/TGFB2/PNPLA7/SP6/NDUFAF6/LAMP1/FOXN2/AHNAK/TXNL1/GSK3B/DSCAML1/IL34/F2RL1/IQUB/CTHRC1/WAC/FAM181A/HERC2/TRPM1/MOBP/BMPER/TECTA/RNF165/IRF2/AKAP9/CELF2/CDC42SE2/ARVCF/SLC39A8/SETD7/FGF10/G3BP1/DST/EYA4/SGK3/PPP1R42/TRMO/SORCS3/CDH13/AFF1/CYCS/PCDH11X/MRPL39/ACTL8/COLGALT2/NIT2/CSMD1/MED12L/MEIKIN/UBXN11/TULP4/MICAL2/THSD4/ABCA8/SLC4A8/GABRG3/TLN2/IGHV270/SPPL2A/ARAP2/TRIQK/FNIP1/PLCL1/COL4A4/C3orf67/FAT1/TMED9/CDKAL1/AGBL1/ANO4/VAT1L/NDUFA12/TRIP4/GOLIM4/ANO3/RSPO4/KAZN/DYSF/FAM240B/C2orf76/ZCCHC10/AICDA/TEX33/DCHS2/STK32B/EXOC4/ST18/PARVB/ST6GAL1/OTOF/NR3C2/SH3RF1/SGK1/MPP3/LAMA1/ASIP/NKAIN2/PLPP4/VSNL1/AP2A2/C11orf65/IL19/SZT2/WDR41/CHN2/INSIG1/DCAF7/SIK1/NCS1/ARHGAP20/SQSTM1/LUZP2/SLC25A48/SESTD1/KRAS/COL25A1/HSPG2/MSH2/RAB1A/DLX5/PDCD1LG2/PPP4R3A/L3MBTL4/ZNF813/COL18A1/MEGF10/MECOM/KCNN2/FARP1/GGACT/PRRC2C/DMXL1/MAPK7/MPZL1/ATXN7L1/NCKAP5/ABCB11/EPC2/CCDC134/BLOC1S4/SH2D4B/CDH1/CEP295NL/PIFO/HMCN1/AHCYL1/SPRED2/C10orf71/D | GWAS Catalog; Nat Genet. 2019 Mar;51(3):404-413. |

| | | |
|---|---|---|
| | ACH1/ELMO1/HECW1/ZAP70/FBXL13/EPOP/TNXB/LMOD3/SLC4A1AP/CCDC85C/SDR9C7/SLC14A2/SYPL1/FAM163A/NRXN1/HDAC9/KCNN3/PDE1A/SORD/RPS20P25/TMEM94/PCSK6/SLMAP/CSNK2A1/PTPRS/SCAPER/UGT1A10/CACNA2D3/LIMS2/CAMK4/PLEKHG1/RELN/FANCD2/RAB20/PDS5B/SPSB1/CDC42EP3/ADCY8/SLC44A5/SLC9A9/TNRC6C/RHBDF1/ELL/CENPM/SFT2D2/KBTBD8/CDON/RBMS3/LRAT/NARS2/BMP1/FBXO40/CASP12/TENM4/FAM19A5/RDX/NAALADL2/JPH3/TBXAS1/HRK/BDH1/ADARB2/GRIN2B/EGLN3/ANTXR1/C9orf92/STK24/DVL2/ERBB4/DAPL1/ANKRD22/AKR7A3/CEP63/DLC1/RASSF8/RAPGEF5/SEC24B/TUSC1/PUM3/TOP1/MYRIP/PUS1/TMEM106B/SENP7/TRIM56/MAP4K4/RNF6/MGME1/LINC02210CRHR1/IRAK1BP1/BICRA/CCDC112/SELENOO/MDGA2/TFEB/DNAH6/LIN28B/C2orf83/PAK2/PCDH7/C5orf64/SLC2A9/PCNX1/POLN/WNT3/OFCC1/GPR180/MRPL45/CFAP74/SELP/USP10/RASSF5/KSR2/PAX2/ABCA1/MAP2K5/ANGPT4/ARIH1/CKAP5/ETS1/PEX6/TMCO4/TAS2R5/NEK10/MSX2/NEGR1/PKNOX2/CPM/CTNND2/AOX1/IQGAP2/PPP5D1/SYNGAP1/ERO1A/GTF2H3/SHANK2/C16orf95/TIAM2/NME9/NRXN3/FAM83E/RIMBP2/STK11/DIP2C/C5orf67/BMP2/CADM2/ADGRL2/SEMA3A/TXN/FAT3/KIFC3/SLC28A1/RBM19/DISC1/EFR3A/UTS2B/MPP7/IRF6/BSG/GAB2/GABRA2/TENT5A/OSTN/EPDR1/HBEGF/IL17RD/PDE7B/ADAMTS1/JCAD/PSMC3/NCR2/SLC10A2/ETF1/IL6/PMAIP1/F2R/CDC5L/FZD6/ACSL6/BANK1/ZNF862/FOXE1/SOBP/HTR7/PPP1R3B/EXOC1L/CDH19/CLDN18/STEAP3/CD24/DDX25/OTOGL/SEPT5/TST/RAPGEF6/ZNF117/CD300LG/TREML2/ATM/HYI/KCNK1/CADPS2/TSPAN16/CDR2L/KIAA0232/TIMP2/STRADA/YAP1/SRCIN1/FRMD4B/HMGA2/LINGO1/ZNF90/TCF15/SEM1/ZNRF4/UGT1A8/LIPC/FANCD2OS/ALDH4A1/MACROD2/RORA/PAX5/CCDC171/ZNF438/THSD7A/NHLRC3/SERPINE1/DIRAS2/OVOL2/BAALC/GDAP1L1/ZNF320/RPL3P8/KLHL36/MLN/LRIG3/PSMC6/LRRIQ3/SERINC5/CCZ1B/C9orf152/CNGB1/SPATA48/RAB3D/ALDH1A2/ZNF468/STEAP1B | |
| Dementia with Lewy body | APOE/SNCA/KRTCAP2/BCL7C/FRMD3/NTS/RASSF9/HPS5/SAA1/MYO7A/HMX3/ACADSB/POT1/C7orf77/PARVB/PDLIM5/CTNND2/VPS36/GARNL3 | GWAS Catalog |
