## Supplementary Table 3 for "Zonation-dependent single-endothelial cell transcriptomic changes in the aged brain"

| Human DEG orthologs | Disease / trait association | GWAS *P*-value | GWAS catalog accession ID / study |
|---|---|---|---|
| NBEAL1 | White matter hyperintensity burden<br>Small-vessel ischemic stroke | $5 \times 10^{-8}$<br>$4 \times 10^{-7}$ | GCST003013<br>GCST005841 |
| SYNE2 | Ischemic stroke | $4.71 \times 10^{-6}$ | Lancet Neurol. 2016 Jun; 15(7): 695–707. |
| VWF | von Willebrand factor levels in ischaemic stroke and hyperhomocysteinaemia | $5 \times 10^{-6}$ | GCST004598 |
| CDKN1A | Ischemic stroke | $2 \times 10^{-7}$ | GCST005843 |
| SWAP70 | Ischemic stroke | $2 \times 10^{-7}$ | GCST005843 |
| SLC50A1 | Parkinson’s disease | $5 \times 10^{-8}$ | GCST001430 |
| ALAS1 | Parkinson’s disease | $3.2 \times 10^{-8}$ | Nat Genet. 2017 Oct; 49(10): 1511–1516. |
| KTN1 | Parkinson’s disease | $2 \times 10^{-7}$ | GCST004902 |
| TPM1 | Parkinson’s disease (age of onset)<br>Alzheimer’s disease | $9 \times 10^{-11}$<br>$3.35 \times 10^{-8}$ | GCST003652<br>Nat Genet. 2019 Mar;51(3):404-413. |
| PAM | Parkinson’s disease | $2 \times 10^{-7}$ | GCST004902 |
| ADAMTS1 | Alzheimer’s disease | $3 \times 10^{-8}$ | GCST007511 |
| DLC1 | Alzheimer’s disease | $4 \times 10^{-6}$ | GCST003815 |
| AHNAK | Age of onset in Alzheimer’s disease<br>Alzheimer’s disease | $9 \times 10^{-8}$<br>$2 \times 10^{-7}$ | GCST003427<br>GCST005549 |
| RDX | Alzheimer’s disease | $3 \times 10^{-6}$ | GCST003815 |
| IRF2 | Hippocampal volume in Alzheimer's disease | $2 \times 10^{-7}$ | GCST006993 |
| CD2AP | Alzheimer’s disease | $5 \times 10^{-11}$ | GCST002245 |
| GOLIM4 | Cerebrospinal fluid total tau levels in Alzheimer's disease | $5 \times 10^{-7}$ | GCST006991 |
| HMCN1 | Alzheimer’s disease | $1 \times 10^{-6}$ | GCST003815 |
| MECOM | Alzheimer’s disease | $9 \times 10^{-7}$ | GCST003815 |
| SIK1 | Age of onset in Alzheimer’s disease | $7 \times 10^{-7}$ | GCST003427 |
| PARVB | Alzheimer’s disease | $5 \times 10^{-7}$ | GCST001915 |
| TGFB2 | Age of onset in Alzheimer’s disease | $8 \times 10^{-8}$ | GCST003427 |
| LIMS2 | Alzheimer’s disease | $2 \times 10^{-6}$ | GCST001915 |
| CD42EP3 | Total ventricular volume in Alzheimer's disease | $2 \times 10^{-6}$ | GCST000892 |
| PRRC2C | Alzheimer’s disease | $9 \times 10^{-7}$ | GCST001915 |
| RORA | Total ventricular volume in Alzheimer's disease | $3 \times 10^{-6}$ | GCST000892 |
| ADGRL2 | Alzheimer’s disease | $9 \times 10^{-6}$ | GCST003815 |
| PAK2 | Total ventricular volume in Alzheimer's disease | $5 \times 10^{-6}$ | GCST000892 |
| KIF1C | Alzheimer’s disease | $9.16 \times 10^{-10}$ | Nat Genet. 2019 Mar;51(3):404-413. |
| MPZL1 | Alzheimer’s disease | $1 \times 10^{-6}$ | GCST001658 |

| | | | |
|---|---|---|---|
| RAPGEF5 | Alzheimer’s disease | $4 \times 10^{-6}$ | GCST003815 |
| LEF1 | Amyotrophic lateral sclerosis | $3 \times 10^{-6}$ | GCST002337 |
| WAPL | Amyotrophic lateral sclerosis | $3 \times 10^{-6}$ | GCST002337 |
| ATXN1 | Amyotrophic lateral sclerosis | $4 \times 10^{-6}$ | GCST000406 |
| ARHGEF2 | Age of onset in amyotrophic lateral sclerosis | $1 \times 10^{-6}$ | GCST001663 |
| EGR1 | Amyotrophic lateral sclerosis | $2 \times 10^{-6}$ | GCST004791 |
| SEM1 | Age on onset in frontotemporal dementia<br>Alzheimer’s disease | $2 \times 10^{-6}$<br>$2 \times 10^{-6}$ | GCST006149<br>GCST003815 |
| RRBP1 | Frontotemporal dementia | $1 \times 10^{-6}$ | GCST006147 |
| XDH | Multiple systems atrophy | $6 \times 10^{-6}$ | GCST003784 |
| ASCC3 | Multiple systems atrophy | $3.66 \times 10^{-6}$ | Neurology. 2016 Oct 11; 87(15): 1591–1598. |
