## Supplementary Table 4 for "Zonation-dependent single-endothelial cell transcriptomic changes in the aged brain"

| Human DEG Orthologs[i] | Concordant with normal aged brain | Concordant with AD brain | Confirmation of expression in human brain endothelium | Zone(s) of protein expression in endothelium[ii] | Other sites of protein expression[iii] |
|---|---|---|---|---|---|
| **ABCB1** | ✓ | | Human Protein Atlas database[iv] showed expression in endothelium by IHC, confirmed by Bendayan et al, 2009. | A, C | None |
| **ACER2** | ✓ | | Human Protein Atlas database showed expression in human cerebral cortex by bulk RT-PCR but IHC data was not available.<br>Allen Human Brain Cell Types snRNA-Seq data[v] showed expression in endothelial cell with low counts. | Not applicable | Not applicable |
| *ADAMTS1* | ✓ | | Human Protein Atlas database showed expression in human cerebral cortex by bulk RT-PCR, but IHC showed absent staining in endothelium.<br>Allen Human Brain Cell Types snRNA-Seq data found *no expression* in endothelial cells<br>Differentially upregulated in human glioma (Martino-Echarri et al, 2014) and sporadic AD (Medoro et al, 2019) but unclear if expressed in endothelium. | Not applicable | Not applicable |
| **AHNAK** | ✓ | | Human Protein Atlas database showed expression in endothelium by IHC. | C, V | Neuronal cell bodies |
| **APOLD1** | ✓ | | Human Protein Atlas database showed no expression in endothelial cells by IHC.<br>Allen Human Brain Cell Types snRNA-Seq data showed expression in endothelial cell with high counts. | Not applicable | Neuronal cell bodies and nuclei |
| **ARHGAP31** | ✓ | | Human Protein Atlas database showed expression in endothelium by IHC. | C | Neuronal cell bodies and neuropils, glial cells |
| *ATOX1* | | ✓ | Human Protein Atlas database showed expression in human cerebral cortex by bulk RT-PCR but IHC data was not available.<br>Allen Human Brain Cell Types snRNA-Seq data did not find expression in endothelial cells.<br>No staining visualized by IHC even though clearly expressed in cerebral cortex by western blot (Davies et al, 2013), localized predominantly to neurons but also some endothelium in a study by Naeve et al, 1999 using in situ hybridization and IHC in rat brain. | Not applicable | Not applicable |
| **CDKN1A** | × | | Human Protein Atlas showed no expression in endothelium by IHC.<br>Allen Human Brain Cell Types snRNA-Seq data showed expression in endothelial cells with low counts. | Not applicable | Neuronal cell bodies (only in certain samples) |

| | | | | | |
|---|---|---|---|---|---|
| | | | Expression might be inducible under certain conditions, predominantly in mitotic and/or tip cells of endothelium in mouse shown by Sabbagh et al, 2018. | | |
| **CLDN5** | ✕ | | Human Protein Atlas database showed expression in endothelium by IHC, confirmed by Du et al, 2017. | A, C, V | None |
| **CXCL12** | ✕ | | Human Protein Atlas database showed expression in endothelium by IHC, confirmed by Krumbholz et al, 2005. | (very weak staining) | Neuronal cell bodies and some neuropils |
| **EPAS1** | ✓ | ✕ | Human Protein Atlas database showed expression in human cerebral cortex by bulk RT-PCR but IHC data was not available.<br>Allen Human Brain Cell Types snRNA-Seq data showed expression in endothelial cells with high counts, confirmed by Grubman et al, 2019 by snRNA-Seq. | Not applicable | Not applicable |
| **ESAM** | | | Human Protein Atlas database showed expression in endothelium by IHC. | A, C, V | Neuropils[vii] |
| **FLT1** | ✓ | | Human Protein Atlas database showed expression in endothelium by IHC, confirmed by Uranishi et al, 2001. | C, V | None[viii] |
| **FOXQ1** | ✕ | | Human Protein Atlas database showed expression in human cerebral cortex by bulk RT-PCR but IHC data was not available.<br>Allen Human Brain Cell Types snRNA-Seq data showed expression in endothelial cells with low counts. | Not applicable | Not applicable |
| *GPR116* | ✓ | | Human Protein Atlas database showed expression in human cerebral cortex by bulk RT-PCR but IHC data was not available.<br>Expression data not available from Allen Human Brain Cell Types snRNA-Seq database.<br>Expression in mouse brain endothelial cells by RT-PCR (Wallgard et al, 2008) and in situ hybridization (Ehrlich et al, 2018). | Not applicable | Not applicable |
| **HES1** | ✓ | | Human Protein Atlas database showed expression in human cerebral cortex by bulk RT-PCR but IHC data was not available.<br>Allen Human Brain Cell Types snRNA-Seq data showed expression in endothelial cells with low counts, confirmed by Murphy et al, 2009 by IF. | Uncertain, only endothelial cells in AVMs shown to express HES1 | None[viii] |
| **ID3** | ✕ | | Human Protein Atlas database showed expression in endothelium by IHC. | C[ix] | Neuronal cell bodies and neuropils, glial cells |
| **IFITM3** | ✕ | ✓ | Human Protein Atlas database showed expression in endothelium by IHC. | C, V | Neuronal cell bodies[x] |
| **IGFBP7** | ✓ | | Human Protein Atlas database showed expression in endothelium by IHC, confirmed by Iqbal U et al, 2010. | (very weak staining) | Neuronal cell bodies |
| **IQGAP1** | ✓ | | Human Protein Atlas database showed expression in endothelium by IHC. | A, C, V | None |

| | | | | | |
|---|---|---|---|---|---|
| **ITGA6** | ✓ | | Human Protein Atlas database showed expression in endothelium by IHC. | (very weak staining)[xi] | Neuropils |
| **KANK3** | ✕ | | Human Protein Atlas database showed expression in endothelium by IHC. | C, V | None |
| **LIMA1** | | ✕ | Human Protein Atlas database showed expression in endothelium by IHC. | A, C | Neuronal cell bodies and neuropils, glial cells |
| **LMO2** | ✓ | | Human Protein Atlas database showed expression in endothelium by IHC, confirmed by Gratzinger et al, 2015. | A, C, V | Neuronal cell bodies and neuropils, glial cells |
| **LY6E** | ✓ | | Human Protein Atlas database showed expression in human cerebral cortex by bulk RT-PCR but IHC data was not available.<br>Allen Human Brain Cell Types snRNA-Seq data showed expression in endothelial cells with low counts. | Not applicable | Not applicable |
| **MECOM** | ✓ | | Human Protein Atlas database showed expression in endothelium by IHC, confirmed by Hou et al, 2016 by IHC. | A, C, V | Neuronal cell bodies and neuropils, glial cells |
| **MFSD2A** | | ✓ | Human Protein Atlas database showed expression in endothelium by IHC. | A, C | Neuronal cell bodies, glial cells, segmental neuropils |
| **MYL12A** | ✕ | | Human Protein Atlas database showed expression in endothelium by IHC. | C | Neuropils |
| **NOSTRIN** | ✕ | ✓ | Human Protein Atlas database showed expression in endothelium by IHC. | C | Neuronal cell bodies and neuropils, glial cells |
| **PLTP** | ✕ | | Annotated by authors of Human Protein Atlas to be absent in brain endothelial cells, but detailed inspection found there might be very faint staining in one of the images.<br>Allen Human Brain Cell Types snRNA-Seq data showed no expression in endothelial cell. See also Vuletic et al, 2003 for expression in human AD brains. | C | Neuronal cell bodies and neuropils |
| **PTPRB** | ✓ | | Annotated by authors of Human Protein Atlas to be absent in brain endothelial cells, difficult to judge by inspection due to intense neuropil staining.<br>Allen Human Brain Cell Types snRNA-Seq data showed expression in endothelial cells with low counts. | Not applicable | Neuropils |

| | | | | | |
|---|---|---|---|---|---|
| **RGCC** | × | | Human Protein Atlas database showed expression in endothelium by IHC. | A, C, V | Neuropils, neuronal cell bodies, glial cells |
| **RHOC** | × | | Human Protein Atlas database showed no expression in endothelium by IHC.<br>Allen Human Brain Cell Types snRNA-Seq data showed expression in endothelial cell with low counts | Not applicable | Not applicable |
| **SDPR** | ✓ | | Human Protein Atlas database showed expression in endothelium by IHC. | C, V | None |
| **SLC2A1** | × | ✓ | Human Protein Atlas database showed expression in endothelium by IHC, confirmed by Cornford et al, 2005. | C | Glial cells[xii] |
| **SLC7A5** | × | | Human Protein Atlas database showed expression in endothelium by IHC. | A, C | Neuronal cell bodies and neuropils |
| **SLC9A3R2** | | × | Human Protein Atlas database showed expression in endothelium by IHC, confirmed by Wallgard et al, 2008. | A, C, V | None[viii] |
| **TAGLN2** | × | | Human Protein Atlas database showed expression in endothelium by IHC. | C | None[viii] |
| **TIMP3** | ✓ | | Human Protein Atlas database showed expression in human cerebral cortex by bulk RT-PCR but IHC data was not available.<br>Allen Human Brain Cell Types snRNA-Seq data showed expression in endothelial cell with high counts. | Not applicable | Not applicable |
| **TINAGL1** | × | | Human Protein Atlas database showed expressed in endothelium by IHC.<br>Allen Human Brain Cell Types snRNA-Seq data showed no expression in endothelial cells. | C | None |
| **TSC22D1** | × | | Human Protein Atlas database showed expression in human cerebral cortex by bulk RT-PCR but IHC data was not available.<br>Allen Human Brain Cell Types snRNA-Seq data showed expression in endothelial cells with intermediate counts. | Not applicable | Not applicable |
| **UACA** | ✓ | | Annotated by authors of Human Protein Atlas to be absent in brain endothelial cells, but detailed inspection found there might be weak staining in some segments of vessel endothelia. | A, C, V[xiii] | Neuronal cell bodies |
| **VWF** | ✓ | | Human Protein Atlas database showed expression in endothelium by IHC, confirmed by Macdonald et al, 2010. | V[xiii] | None[viii] |

Abbreviations and annotations: A: arteriole, AD: Alzheimer's disease; C: capillary, DEG: differentially expressed genes, IF: immunofluorescence; IHC: immunohistochemistry, RT-PCR: reverse transcription-polymerase chain reaction, scRNA-Seq: single cell RNA sequencing; snRNA-Seq: single nucleus RNA sequencing; V: venule; genes in bold font: have evidence of human brain endothelial expression.

Footnotes

i Gene names were chosen in accordance with Human Protein Atlas. Bold: confirmed expression in endothelium. Italic: unable to find evidence for endothelial expression.

ii Annotated by inspecting available human brain images from Human Protein Atlas and literature, may not be accurate due to sampling error.

iii Annotated by inspecting available human brain images from Human Protein Atlas and literature, cross-compared with Lee et al 2017, may not be accurate due to sampling error.
iv Retrieved from https://www.proteinatlas.org/humanproteome/tissue/brain, on 10/8/2019
v Retrieved from https://celltypes.brain-map.org/rnaseq/human on 10/8/2019, data was based on snRNA-Seq of cells from the middle temporal gyrus.
vi Difficult to determine due to weak nuclear staining.
vii Lee et al 2017 annotated it as specific for endothelium, we disagree as neuropil staining was clearly demonstrated in all samples in Human Protein Atlas.
viii Not commented by Lee et al to be specific for endothelium.
ix Difficult to determine due to weak nuclear staining, not present in all samples.
x Lee et al 2017 annotated it as specific for endothelium, we disagree as neuronal cell bodies were clearly stained in one sample in Human Protein Atlas and is unlikely due lipofuscin staining.
xi Difficult to determine due to intense neuropil staining.
xii Lee et al 2017 annotated it as specific for endothelium by IHC, however a transmission electron microscopy study by Conford et al 2005.
xiii Only in some segments of vessels.
